# Mdm38/LETM1 couples ion homeostasis and proteostatic mechanisms in the inner mitochondrial membrane

**DOI:** 10.1101/2025.01.30.635785

**Authors:** Iryna Bohovych, Gabriella Menezes da Silva, Saieeda Fabia Ali, Emma J. Bergmeyer, Edward M. Germany, Adarsh Mayank, James A. Wohlschlegel, Jason C. Casler, Muhammad Arifur Rahman, Taras Y. Nazarko, Maureen Tarsio, Takuya Shiota, Laura L. Lackner, Steven M. Claypool, Patricia M. Kane, Antoni Barrientos, Oleh Khalimonchuk

## Abstract

The mitochondrial inner membrane is among the most protein-dense cellular membranes. Its functional integrity is maintained through a concerted action of several conserved mechanisms that are far from clear. Here, using the baker’s yeast model, we functionally characterize Mdm38/LETM1, a disease-related protein implicated in mitochondrial translation and ion homeostasis, although the molecular basis of these connections remains elusive. Our findings reveal a novel role for Mdm38 in maintaining protein homeostasis within the inner membrane. Specifically, we demonstrate that Mdm38 is required for mitochondrial iron homeostasis and for signaling iron bioavailability from mitochondria to vacuoles. These processes are linked to the m-AAA quality control protease, whose unrestrained activity disrupts the assembly and stability of respiratory chain complexes in Mdm38-deficient cells. Our study highlights the central role of Mdm38 in mitochondrial biology and reveals how it couples proteostatic mechanisms to ion homeostasis across subcellular compartments.

## INTRODUCTION

Mitochondria are essential organelles that serve as multifaceted hubs for cellular bioenergetics, metabolism, and signaling [1–3]. These functions are critical for cellular physiology, and disruptions in mitochondrial activities are linked to over 150 human disorders, including both hereditary and age-related diseases [4,5]. Mitochondria are enclosed by two membranes: the outer (OMM) and inner (IMM) mitochondrial membranes, the latter being one of the most protein-rich cellular membranes, housing more than half of the mitochondrion’s ∼1,300 proteins[6,7]. Many of these proteins form the complexes of the oxidative phosphorylation system (OXPHOS), which generates cellular ATP. A unique feature of the OXPHOS complexes is their mosaic composition, comprising subunits encoded by both nuclear DNA –synthesized at cytosolic ribosomes and imported into the mitochondria– and mitochondrial DNA (mtDNA), which are essential core subunits synthesized by mitochondrial ribosomes and inserted co-translationally into the IMM[5,8,9].

Studies in budding yeast have identified a handful of proteins that facilitate the co-translational membrane insertion of mtDNA-encoded OXPHOS subunits into the IMM. These include the evolutionarily conserved membrane insertase Oxa1 and auxiliary factors like Mba1 and Mdm38 (also known as LETM1 in higher eukaryotes) [8,10]. Mdm38 is a IMM-anchored protein proposed to perform a variety of functions, such as ribosome tethering to IMM and regulation of mitochondrial translation [11–14] as well as control of mitochondrial volume through regulation of ion homeostasis[15–20]. However, the precise roles of Mdm38 remain poorly defined, and some of its proposed functions are still debated, highlighting the need for further research to clarify its contribution to mitochondrial biology.

The IMM proteome is highly susceptible to disruptions, such as protein misfolding, aggregation, or improper assembly, which can compromise mitochondrial function and lead to proteostatic imbalances. Such disruptions result in metabolic and ion imbalance, bioenergetic deficit, and increased oxidative damage. To avert such unwanted scenarios, mitochondria rely on several conserved mechanisms to preserve the integrity of IMM resident proteins. One such mechanism is represented by IMM proteases like the ATP-fueled proteolytic complexes called i-AAA (intermembrane space, IMS-facing AAA protease, Yme1) and m-AAA (matrix-facing AAA protease comprised of Yta10 and Yta12 subunits), alongside ATP-independent proteases such as Oma1 and Pcp1[21,22]. Not only do these conserved proteolytic modules survey and maintain the IMM proteome, but they also play roles in shaping the molecular architecture of the membrane. In particular, the m-AAA protease has dual functions in proteolysis and chaperoning for IMM proteins and matrix-localized proteins that are adjacent to the IMM, such as the bL32m subunit of the mitochondrial ribosome[23,24]. Similarly, Oma1, initially identified as an accessory protease to the m-AAA, has emerged as a critical regulator of cellular stress responses and signaling in both yeast and mammalian cells[21,22]. However, despite recent advances, much remains to be uncovered about their operational mechanisms, interactions within the IMM proteome, and regulatory controls thereof.

Here, we used the budding yeast model to address gaps in our understanding of Mdm38’s role in mitochondrial biology. We demonstrate that the loss of Mdm38 function disrupts IMM proteostasis, leading to altered tolerance to cobalt ions and impaired iron bioavailability. Such disruption activates the m-AAA protease, causing the rapid degradation of newly synthesized core OXPHOS subunits. Our findings reveal the role of Mdm38/LETM1 in coupling proteostatic mechanisms with ion homeostasis across subcellular compartments, and highlight how unrestrained m-AAA protease activity exacerbates mitochondrial dysfunction in the absence of Mdm38, underscoring its importance in maintaining mitochondrial integrity.

## RESULTS

### Specific ion homeostasis-perturbing compounds activate IMM protein quality control mechanisms

Previous findings by our group and others have established that compounds such as the protonophore Carbonyl Cyanide m-ChloroPhenylhydrazone (CCCP), an OXPHOS uncoupler, can activate mitochondrial protein quality control mechanisms in the IMM, particularly involving the stress-sensitive protease Oma1[25–27]. To further explore how other ion homeostasis-perturbing compounds affect IMM proteostasis, we sought to examine the impact of several such compounds on Oma1 activity. To that end, we isolated mitochondria from wild-type (WT) cells expressing Oma1 tagged with a benign C-terminal 13xMyc epitope tag and subjected them to acute treatments with CCCP (used as a positive control), the K^+^/H^+^ ionophore nigericin, NaCl, and KCl. We then analyzed the behavior of the oligomeric Oma1 complex using blue-native gel electrophoresis (BN-PAGE). In yeast, Oma1 activation results in the formation of a labile, high-mass Oma1 complex that is destabilized under BN-PAGE conditions, whereas steady-state levels of Oma1 remain unaffected[25,28]. As expected, mitochondria from CCCP-treated cells showed a marked destabilization of Oma1 oligomers compared to mock-treated cells (Fig. 1A). Notably, Oma1 oligomers were also destabilized in mitochondrial lysates from cells acutely treated with K^+^/H^+^ ionophore nigericin, whereas Oma1 steady-state levels were unaltered (Fig. 1A). In contrast, treatment with NaCl or KCl did not have any appreciable effect on the Oma1 complex stability (data not shown). These results suggest that nigericin may trigger IMM protein quality control mechanisms that involve Oma1, likely via perturbation of ion homeostasis.

**Figure 1.**
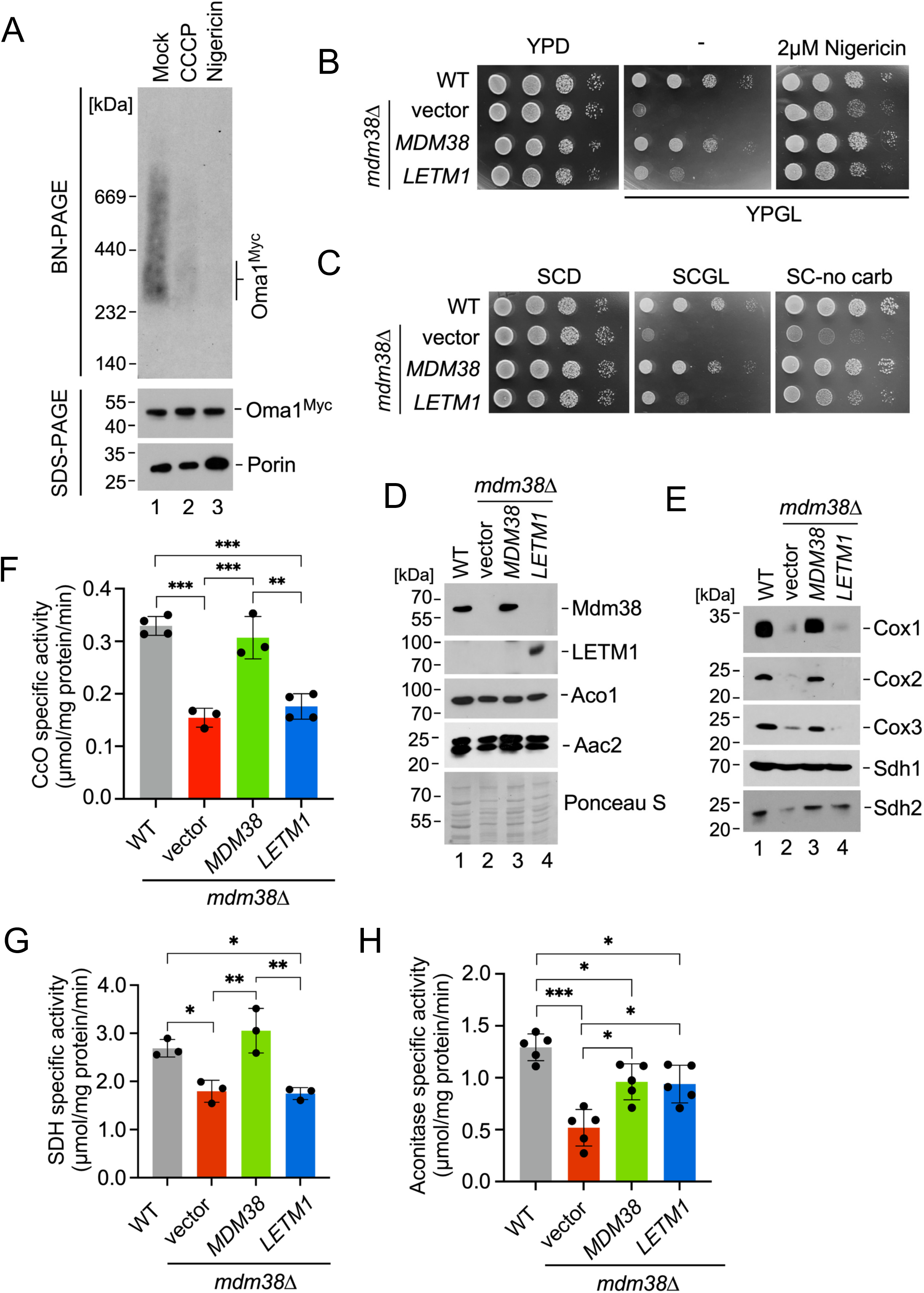
Loss of Mdm38 impacts OXPHOS and other IMM-linked mitochondrial processes. (A) Blue-native (BN)-PAGE analysis of Oma1 high-mass complexes in mitochondria from chromosomally tagged Oma1-13xMyc cells that were mock-treated or challenged for two hours with an uncoupler CCCP or a K^+^/H^+^ antiporter nigericin, lysed with digitonin. Coomassie G250- stained membrane shows equal amounts (100 µg) of total mitochondria were analyzed. The bottom panel shows an immunoblot of SDS-PAGE analysis used to visualize steady-state levels of the individual proteins detected with indicated antibodies. (B) Growth assays of indicated yeast cells on rich medium supplemented with dextrose (YPD) or glycerol/lactate mix (YPGL) as the carbon source, with or without nigericin. (C) Serial growth assays of the wild-type (WT) yeast strain and the *mdm38*Δ mutant containing empty vector or transformed with low-copy plasmids harboring *MDM38* or its human ortholog *LETM1*. Cells were spotted onto synthetic complete medium supplemented with dextrose (SCD) or glycerol/lactate mix (SCGL) as the carbon source or without supplementation (SC-no carb). (D) Immunoblot analysis showing steady-state levels of indicated proteins in mitochondria from the cells described above. Ponceau S staining was used as a control for protein loading. (E) Steady-state levels of individual proteins representing select OXPHOS complexes. Cox1, Cox2, and Cox3 are the mtDNA-encoded subunits of complex IV; Sdh1 and Sdh2 are nucleus- encoded subunits of complex II. (F-H) Specific enzymatic activities of complex IV (cytochrome *c* oxidase, CcO; panel F), complex II (succinate dehydrogenase, SDH; panel G), and aconitase (panel H) in mitochondrial lysates from indicated cells. Bars show mean ± SD; n=3-5 biological replicates. Asterisks indicate a statistically significant difference by one-way ANOVA test (*p<0.05, **p<0.01, ***p<0.001).

Nigericin, functionally a K^+^/H^+^ antiporter, is known to exert specific effects in budding yeast, particularly in relation to Mdm38. Previous studies have demonstrated that low doses of nigericin can partially restore respiratory function in *mdm38*Δ mutant cells [11,19] (Fig. 1B). To further dissect the functional connection between IMM protein quality control and Mdm38, we first assessed the effect of nigericin on yeast cells. Consistent with previous reports, low doses of nigericin mitigated the respiratory defect in the *mdm38*Δ cells (Fig. 1B). We confirmed that the respiratory competency of the *mdm38*Δ mutant was fully rescued upon re-expression of *MDM38* as indicated by the restored respiratory growth and steady-state levels of cytochrome *c* oxidase (CcO) core subunits Cox1, Cox2, and Cox3 and succinate dehydrogenase (SDH) subunit Sdh2. Interestingly, expressing the human ortholog of Mdm38, *LETM1*, only provided a minor rescue effect in the *mdm38Δ* mutant – likely due to substantial evolutionary divergence between the two proteins [14,20] (Fig. 1B-E).

Biochemical analyses of mitochondria isolated from these strains and controls revealed several key insights: (i) mitochondria isolated from *mdm38*Δ cells cultured for prolonged periods (16 h versus 48 h) retained considerable residual CcO activity, suggesting that Mdm38 is dispensable under these conditions (Fig. 1F and Fig. S1A-S1C); and (ii) beyond CcO, Mdm38 deletion affected the activity and stability of the Krebs cycle enzymes succinate dehydrogenase (SDH) and aconitase (Fig. 1G, 1H). Of note, the activities of ROS scavenging superoxide dismutases Sod1 and Sod2 did not increase in the *mdm38*Δ mutant (Fig. S1D, S1E), indicating that reduced aconitase activity was likely unrelated to oxidative damage. Moreover, the *mdm38*Δ cells exhibited marked resistance to nigericin concentrations higher than wild-type cells (Fig. S1F), although this feature appears to be shared with other mutants deleted for genes encoding key subunits of OXPHOS complexes (data not shown). These results indicate that the consequences of Mdm38 loss extend beyond just OXPHOS complexes harboring mtDNA-encoded subunits, affecting broader aspects of mitochondrial function.

### Functional consequences of Mdm38 loss

To better understand the Mdm38-dependent cellular processes that contribute to IMM proteostasis and mitochondrial respiration, we systematically examined the molecular consequences of Mdm38 deletion. Guided by previous studies [11,14], we first interrogated the connection between Mdm38 and mitochondrial translation using the *cox1*Δ::*ARG8* genetic reporter strain. In this strain, a variant of acetylornithine aminotransferase Arg8 is expressed from the *COX1* locus that encodes for Cox1 core subunit of CcO [29] (Fig. 2A). This experimental setup allows for *in vivo* monitoring of mitochondrial transcriptional and translational activity by assessing the growth of the mutant in medium lacking arginine. We found that the deletion of *MDM38* in the *cox1*Δ::*ARG8* strain had a marginal effect on arginine-dependent growth, indicating that Mdm38 is not essential for mitochondrial transcription and translation (Fig. 2B, 2C). Consistent with this finding, the introduction of the *mdm38*Δ mutation into a mtDNA-intronless background yielded a strain that was phenotypically indistinguishable from the *mdm38*Δ mutant in the intron-containing strain (Fig. S2A). These observations are consistent with previous reports that Mdm38 is not required for normal mitochondrial ribosome assembly or IMM tethering [30,31]. In addition, *in organello* pulse-chase labeling of mitochondrial translational products in intact mitochondria isolated from WT and *mdm38*Δ cells revealed reduced levels of translation products in the latter organelles (Fig. 2D). Together, these data suggest that although Mdm38 may contribute to an increase in the rate of mitochondrial translation, the more significant effect is the rapid turnover of newly synthesized polypeptides in the *mdm38*Δ mutant.

**Figure 2.**
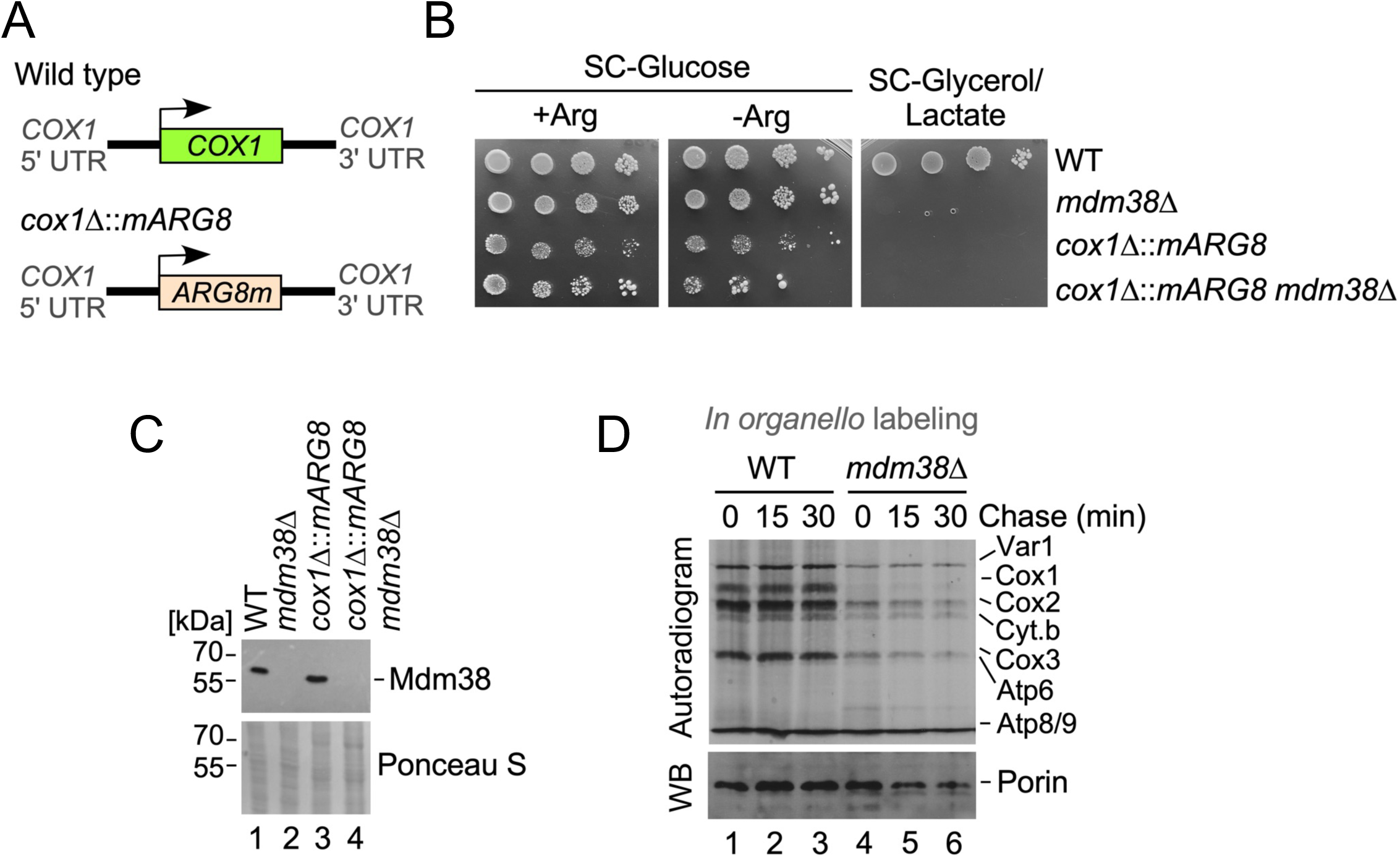
Mdm38 is required for post-translational stability of the mitochondria-borne polypeptides. (A) Schematic of the *cox1*Δ*::mARG8* genetic reporter, compared to wild-type *COX1* locus. (B) Growth assays assessing transcription and translation of the *COX1* ORF from the mitochondrial genome in WT and *mdm38*Δ cells with normal *COX1* and *cox1*Δ*::mARG8* loci on SC-glucose ± arginine or SC-glycerol/lactate. Cells were serially diluted, spotted on respective media, and cultured for 3 (SC-glucose) or 5 (SC- glycerol/lactate) days at 28°C. (C) Steady-state levels of Mdm38 in the mitochondria from cells described in panel B. Ponceau S protein staining was as a loading control. (D) Metabolic pulse-chase labeling of mitochondrial translation products in the intact organelles isolated from wild-type and *mdm38*Δ cells. The autoradiogram shows steady-state levels of the indicated translational products labeled with ^35^S-methionine. The bottom panel shows immunoblot visualizing mitochondrial protein porin, which served as a loading control.

Mdm38-deficient cells have been shown to exhibit mitochondrial network and ultrastructure abnormalities [15,19]. To further explore the impact of Mdm38 deletion on mitochondrial membrane biology, we conducted phospholipid labeling analyses. These experiments revealed no appreciable changes in the steady-state levels of mitochondrial phospholipids in the *mdm38*Δ mutant (Fig. S2B, S2C), indicating the observed defects are not due to altered phospholipid composition of the mitochondrial membranes.

Given that combined deletion of the OM division factor Dnm1 and the IM pro-fusion protein Mgm1 stabilizes the mitochondrial network and permits respiratory growth [32] (Fig. S3A, S3B), we hypothesized that a similar stabilization in Mdm38-deficient cells could potentially rescue their respiratory and other defects. However, in contrast to the *dnm1*Δ *mgm*1Δ strain, the triple *mdm38*Δ *dnm1*Δ *mgm*1Δ mutant remained respiratory deficient Fig. S3B), suggesting that the mitochondrial network defect in the Mdm38-deficient cells is a secondary phenotype and not the root cause of their respiratory deficit.

Interestingly, we observed that the ER-mitochondria and ER-vacuole contact sites, crucial for interorganellar communication and function, were unaltered in the *mdm38*Δ cells (Fig. S3C- F). Also, while Mdm38-deficient cells have been reported to have decreased mitochondrial membrane potential, they exhibit normal mitochondrial protein import *in vitro* when isolated organelles are fully energized [14] (Fig. S4A). To rule out potential *in vivo* import defects, we used Oxa1-Ura3 genetic reporter [33] (Fig. S4B), which consistently did not reveal appreciable changes in protein import across mitochondrial membranes (Fig. S4C).

Altogether, these findings suggest that the mitochondrial respiratory defects in *mdm38***Δ** cells are not primarily linked to alterations in mitochondrial membrane composition, network morphology, or organelle contact sites, but rather to more fundamental disruptions in IMM protein homeostasis and mitochondrial ion metabolism.

### Loss of Mdm38 triggers proteostatic response in the IMM

To gain a more comprehensive insight into the mechanisms underlying Mdm38 deficiency, we used a multiomics approach. First, we explored the metabolic consequences of Mdm38 loss by performing gas chromatography (GC)-MS-based metabolomics analyses of WT and *mdm38*Δ cells. Our analyses revealed significant alterations in several classes of metabolites (Fig. 3A, 3B), including elevated levels of lactate, citrate, xanthine, and aspartate (Fig. 3B). These metabolic changes are consistent with the observed OXPHOS deficits in the *mdm38*Δ mutant but also indicate more specific perturbations in key metabolic pathways. Notably, the accumulation of citrate and altered levels of tricarboxylic acid (TCA) cycle intermediates point to perturbations in the TCA cycle, which aligns with our earlier findings of reduced SDH levels and attenuated activity of aconitase (see Fig. 1G, 1H).

**Figure 3.**
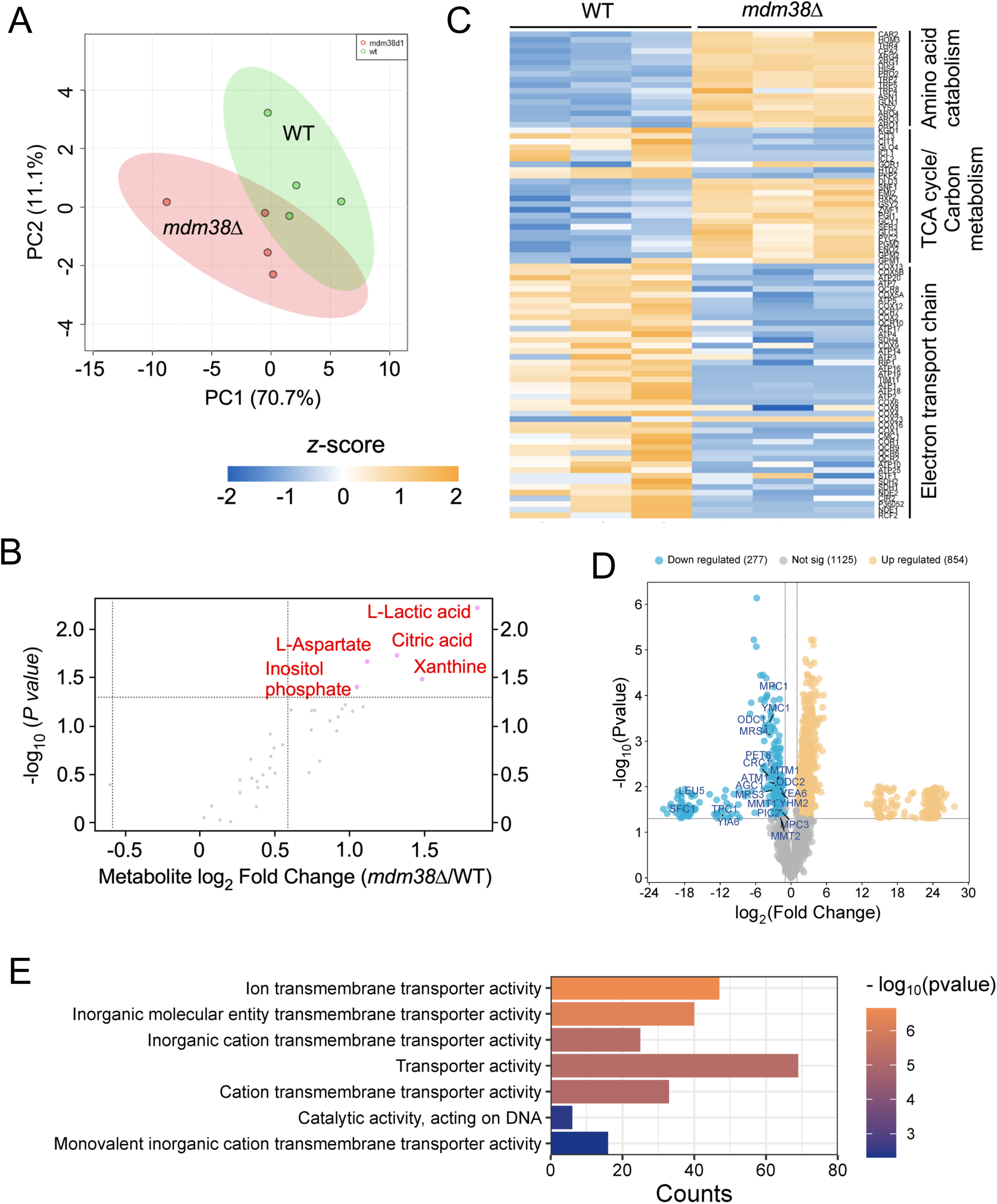
Multi-omics analysis identifies metabolic perturbations and altered mitochondrial protein homeostasis in the Mdm38-deficient cells. (A) Principal component analysis of metabolomic profiles control in wild-type and *mdm38*Δ mutant cells. (B) Volcano plot showing metabolic changes in the *mdm38*Δ mutant cells relative to WT control. The pink circles denote the most altered metabolites with a false discovery rate (FDR) < 0.05. (C) Heat map showing abundance of proteins from quantitative proteomics analysis carried out on mitochondria isolated from log-phase WT and *mdm38*Δ cells. Blue, downregulated; yellow, upregulated. Maps are clustered by indicated mitochondrial processes. (D) Volcano plot showing proteomic changes in wild-type and *mdm38*Δ cells. The blue circles depict downregulated proteins in the *mdm38*Δ mutant with altered IMM transporters highlighted. FDR< 0.05. (E) Protein set enrichment analysis comparing wild-type and *mdm38*Δ cells. See also Supplemental Figure S5.

We next performed quantitative proteomic analysis to gain an unbiased evaluation of proteome changes in Mdm38-deficient cells. As expected, we observed a decrease in the abundance of several OXPHOS proteins and a subset of TCA cycle enzymes. More strikingly, we also detected an increase in proteins associated with amino acid catabolism (Fig. 3C, 3D, and Fig. S5A-C), suggesting a compensatory shift in metabolic pathways in response to Mdm38 loss. Furthermore, we identified significant changes in the levels of several IMM ion and metabolite transporters, including mitoferrin Mrs3, a key protein involved in mitochondrial iron import (Fig. 3D, 3E, and Fig. S5D). This finding is particularly relevant, as it ties into the observed iron homeostasis disruption in *mdm38***Δ** cells, further supporting the idea that Mdm38 may play a role in regulating mitochondrial ion transport and maintaining proper ion homeostasis.

Additionally, we observed a significant upregulation of proteostatic factors in the *mdm38***Δ** mutant, many of which have been previously associated with mitochondrial precursor overaccumulation stress (mPOS) [34,35] (Fig. 4A, 4B). In line with this notion, inducible expression of the mouse dihydrofolate reductase (DHFR) variants [36,37] targeted specifically to the IMS side of the IMM, but not the mitochondrial matrix, further potentiated growth defects of the *mdm38*Δ mutant and not the WT control cells when grown on galactose or glycerol/lactate containing media (Fig. 4C, 4D). This suggests that Mdm38 loss activates a proteostatic response within the IMM, likely as a protective mechanism to cope with the stress induced by improper co-translational insertion of mitochondria-borne proteins, leading to their misfolding and/or aggregation.

**Figure 4.**
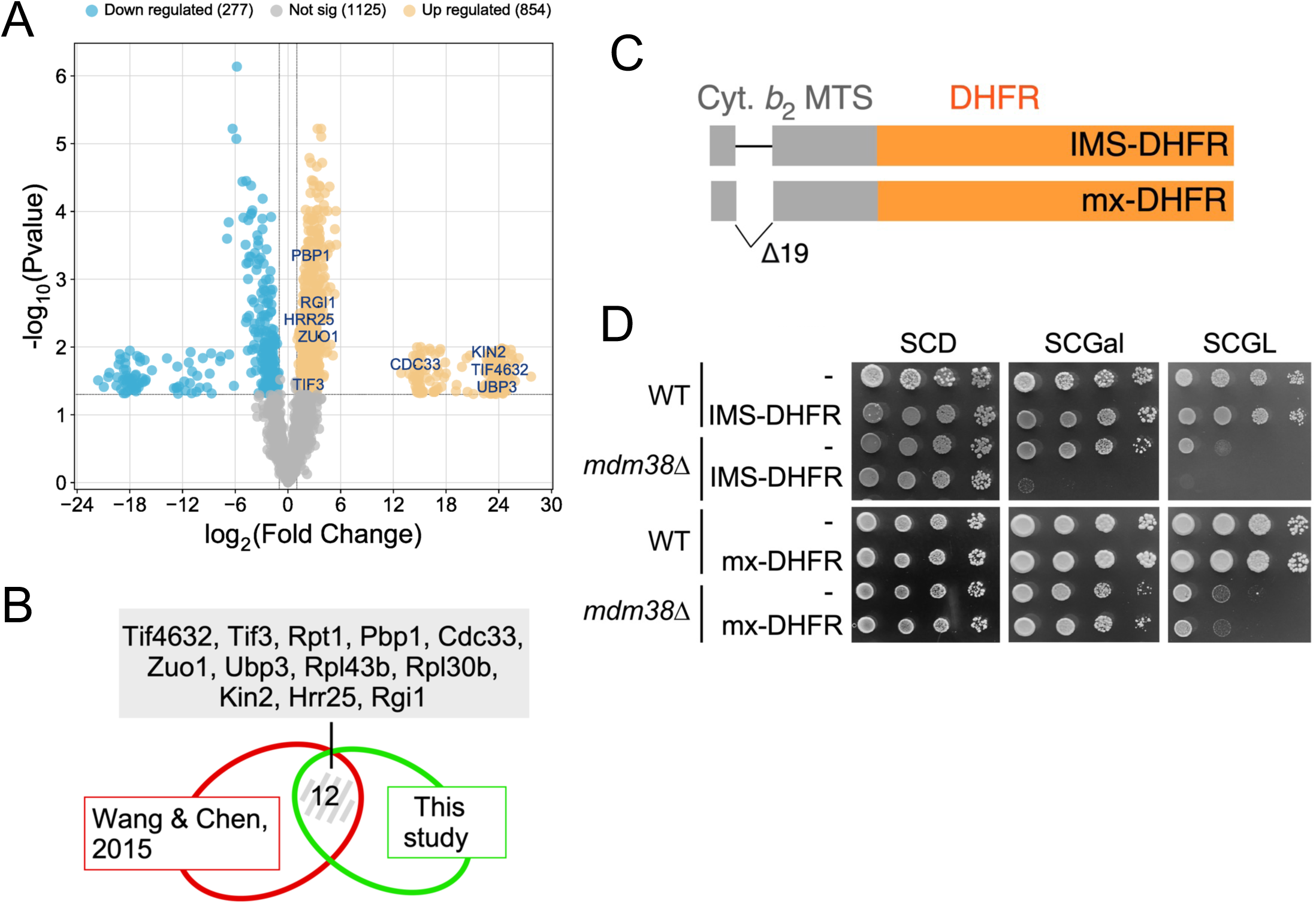
Cells devoid of Mdm38 exhibit signs of proteostatic distress. (A) Volcano plot showing proteomic changes in wild-type and *mdm38*Δ cells. The yellow circles depict upregulated proteins in the *mdm38*Δ mutant with FDR< 0.05. (B) Venn diagram comparing proteomic enrichment of proteostasis-related factors identified in the *mdm38*Δ cells by the present study and mitochondrial precursor overaccumulation stress (mPOS) signature identified by Wang and Chen study[34]. (C) Schematic depiction of galactose-inducible variants of IMS- and matrix-targeted dihydrofolate reductase (DHFR) variants. These constructs are designed to create additional proteostatic pressure in the mitochondrial compartment they are targeted to. (D) Growth assays of indicated yeast cells expressing IMS- and matrix-DHFR on synthetic medium supplemented with dextrose (SCD), galactose (SCGal) or glycerol/lactate mix (SCGL) as the carbon source.

Collectively, these data support a model in which Mdm38 plays a crucial role in maintaining IMM proteostasis and ion homeostasis. Its loss triggers compensatory proteostatic responses, which, while aimed at mitigating stress, ultimately compromise overall mitochondrial function.

### Iron homeostasis is altered in Mdm38-deficient cells

Building on our findings that Mdm38-deleted cells exhibit reduced SDH and aconitase activities, and a previous report that mitoferrin Mrs3 can act as a partial high-copy suppressor of the respiratory deficient phenotype in *mdm38*Δ cells [38], we investigated the relationship between Mdm38 and iron homeostasis. To interrogate this connection, we first assessed the functionality of mitochondrial iron-sulfur cluster biosynthetic machinery in the *mdm38*Δ mutant. A useful readout for this process is mitochondrial lipoylation, as the attachment of lipoyl groups to the pyruvate dehydrogenase subunit dihydrolipoamide acetyltransferase (Lat1) and the dihydrolipoyl succinyltransferase (Kdg2) depends on iron-sulfur cluster-dependent mitochondrial acyl carrier protein (Acp1) [39]. Our analyses revealed that lipoylated protein levels were only partially decreased in the absence of Mdm38, ruling out a gross defect in mitochondrial iron-sulfur cluster biosynthesis (Fig. S6A). This finding is consistent with our earlier observation that heme biosynthesis and trafficking are largely unperturbed in the *mdm38*Δ mutant [40].

Next, we examined the growth of Mdm38-deleted cells under conditions known to affect overall iron homeostasis. We observed that, while the growth of *mdm38*Δ cells was not significantly altered on plates containing the iron chelator bathophenanthrolinedisulfonate (BPS) or supplemented with exogenous iron (ferrous ammonium sulfate) (Fig. S6B), there was a marked resistance to the V-ATPase inhibitor concanamycin A (Conc. A), which compromises vacuolar acidification, thereby affecting iron bioavailability [41] (Fig. 5A). Such resistance is indicative of a defect in iron mobilization due to altered vacuolar pH and suggests that the *mdm38*Δ mutation impacts the fine balance of iron homeostasis within the cell.

**Figure 5.**
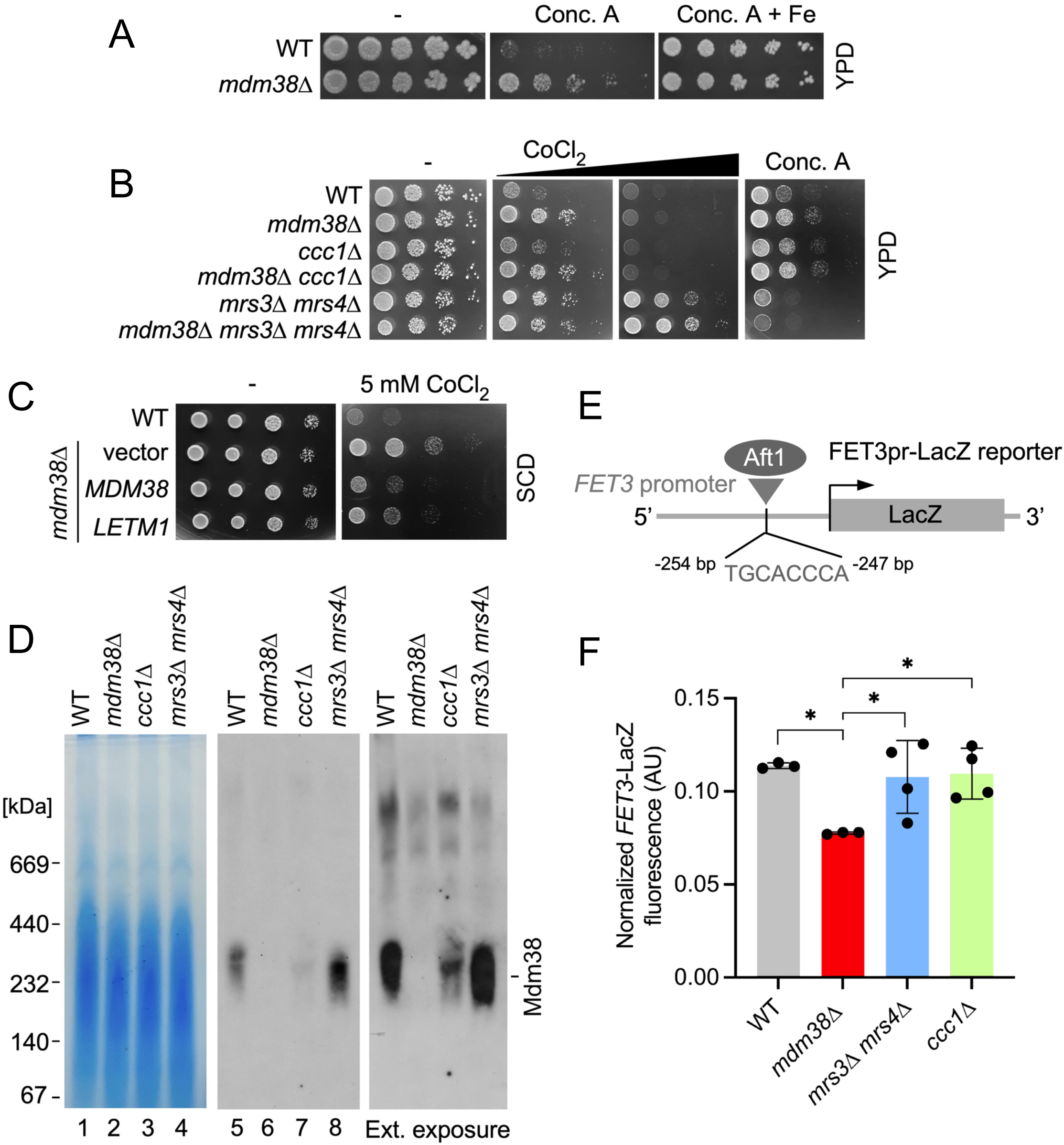
Iron bioavailability is altered in the Mdm38-deficient cells. (A) Growth assays of indicated yeast strains on rich medium with glucose as a carbon source (YPD) with or without 100 nM V-ATPase inhibitor concanamycin A (Conc.A) ± ferrous ammonium sulfate (Fe). (B) Growth of indicated yeast strains on YPD medium ± 100 nM Conc.A or ± lower (4 mM) or higher (6mM) cobalt chloride (CoCl2). (C) Growth assays of indicated yeast cells on synthetic medium supplemented with dextrose (SCD) with or without CoCl2. (D) BN-PAGE analysis of Mdm38 protein complexes in mitochondria from indicated strains. Coomassie G250-stained membrane shows equal amounts (100 µg) of total mitochondria were lysed with digitonin and loaded in each lane. See also Supplemental Figure S6E. (E) Schematic depicting the Aft1-driven transcriptional reporter used in this study. (F) Fluorescence detection analysis showing LacZ reporter expression from the Aft1-regulated *FET3* promoter in the indicated strains. Bars show mean ± SD; n=3-4 biological replicates. Asterisks indicate a statistically significant difference by one-way ANOVA test (*p<0.05).

Consistent with this notion, the WT strain was able to grow on Conc. A-containing plates when supplemented with exogenous iron, whereas no such effect was observed for the *mdm38*Δ cells (Fig. 5A). Likewise, the Mdm38-deficient strain displayed specific marked tolerance to cobalt chloride compared to WT cells (Fig. 5B, 5C). No altered sensitivity of the *mdm38*Δ strain to other mono- or divalent metals was observed (Fig. S6C). This is particularly notable as cobalt metabolism in yeast is tightly linked to iron bioavailability [42]. We confirmed that these phenotypes were indeed related to the absence of Mdm38, as the re-expression of *MDM38* rescued these effects (Fig. 5C). Moreover, growth profiling of canonical OXPHOS mutants further indicated the specificity of these phenotypes (Fig. S6D).

By contrast, and congruent with a model linking Mdm38 to iron homeostasis, cells lacking iron-transporting mitoferrins Mrs3 and Mrs4 also displayed resistance to cobalt chloride, whereas a strain lacking the vacuolar Fe^2+/^Mn^2+^ importer Ccc1 did not (Fig. 5B). This suggests a functional link between Mdm38 and mitochondrial iron transport. Deleting *MDM38* in the *mrs3*Δ *mrs4*Δ genetic background did not result in additive effects, suggesting that Mdm38 and mitoferrins likely function in the same pathway (Fig. 5B). Of note, the *mrs3*Δ *mrs4*Δ double mutant cells did not show appreciable resistance to Conc. A, indicating this phenotype may be more specific to *MDM38* deletion.

BN-PAGE analysis of the Mdm38 high-mass complex in mitochondrial lysates from the WT, *ccc1*Δ and *mrs3*Δ *mrs4*Δ stains revealed a unique trend. In particular, the levels of Mdm38 oligomers inversely responded to mitochondrial iron status in *ccc1*Δ and *mrs3*Δ *mrs4*Δ mutants, where they were reduced and elevated, respectively, whereas the steady-state levels of Mdm38 remained unchanged across the strains (Fig. 5D and Fig. S6E). Concomitant-induced expression of mitoferrins Mrs3 and Mrs4 in the *mdm38*Δ mutant cells partially restored their respiratory capacity (Fig. S6F), further underscoring the functional link between Mdm38 and these iron-transporting proteins.

To investigate the impact of Mdm38 loss on cellular iron metabolism further, we analyzed the activity of a genetic *FET3*pr-LacZ transcriptional reporter composed of the promoter region of the high-affinity iron uptake system component, ferrooxidoreductase Fet3 and LacZ moieties [43] (Fig. 5E). The *FET3* promoter activity is controlled by Aft1- the master regulator of iron metabolism that translocates to the nucleus upon iron deprivation, thereby activating a set of iron-responsive genes called the iron regulon [43–45]. Quantitative fluorescence analysis revealed that in the *mdm38*Δ mutant, Aft1-dependent LacZ expression was significantly diminished compared to WT, *mrs3*Δ *mrs4*Δ, and *ccc1*Δ strains, suggesting altered iron signaling in the absence of Mdm38 (Fig. 5F). In line with this, the expression of the iron-sensitive *HMX1* gene was markedly elevated in the *mdm38*Δ mutant relative to WT (Fig. S6G). The measurements of total intracellular iron levels using inductively coupled plasma mass spectrometry (ICP-MS) showed that *mdm38*Δ mutant cells maintained iron levels similar to WT (Fig. S6H), indicating that total cellular iron uptake remains unaffected. However, mitochondrial iron levels showed a downward trend, albeit not statistically significant (Fig. S6I, S6J). This further suggests that while total iron levels are largely maintained, the distribution of iron between cellular compartments, specifically the mitochondria, is altered in the *mdm38***Δ** mutant.

Collectively, these results suggest a functional connection between Mdm38 and iron homeostasis, particularly in relation to mitochondrial iron transport and bioavailability. The perturbations in mitochondrial iron levels, coupled with changes in iron-responsive gene expression, support a model in which Mdm38 plays a key role in maintaining mitochondrial iron balance. Yet, such iron-related perturbations alone may not fully explain the observed respiratory deficiencies in *mdm38*Δ mutant cells. Instead, these changes likely add another layer to the complex molecular landscape revealed through our proteomic analysis, where ion homeostasis and proteostatic stress further exacerbate mitochondrial dysfunction.

### Loss of Mdm38 has a moderate effect on vacuolar function

Given previous reports that loss of Mdm38 function can induce mitophagy and alter vacuole morphology [19,38], and that the activity of vacuolar V-ATPase is important for maintaining iron homeostasis and mitochondrial respiration [41,46], we further investigated vacuolar function in the Mdm38-deficient cells. The rationale for these studies was further supported by the observation that *mdm38***Δ** mutants exhibit resistance to Conc. A, a V-ATPase inhibitor (Fig. 5A), suggesting a potential underlying defect in V-ATPase function.

Cells lacking V-ATPase activity (*vma2Δ* and *vma3Δ*) are unable to grow at elevated pH, a phenotype that is further exacerbated by the presence of CaCl2 (Fig. 6A). However, there is no appreciable growth impairment in the *mdm38*Δ strain under these conditions, even at high temperatures (Fig. 6A). Further examination of V-ATPase subunit levels and activity in vacuoles isolated from WT and *mdm38*Δ strains revealed a ∼45% decrease in activity compared to WT (Fig. 6B-D).

**Figure 6.**
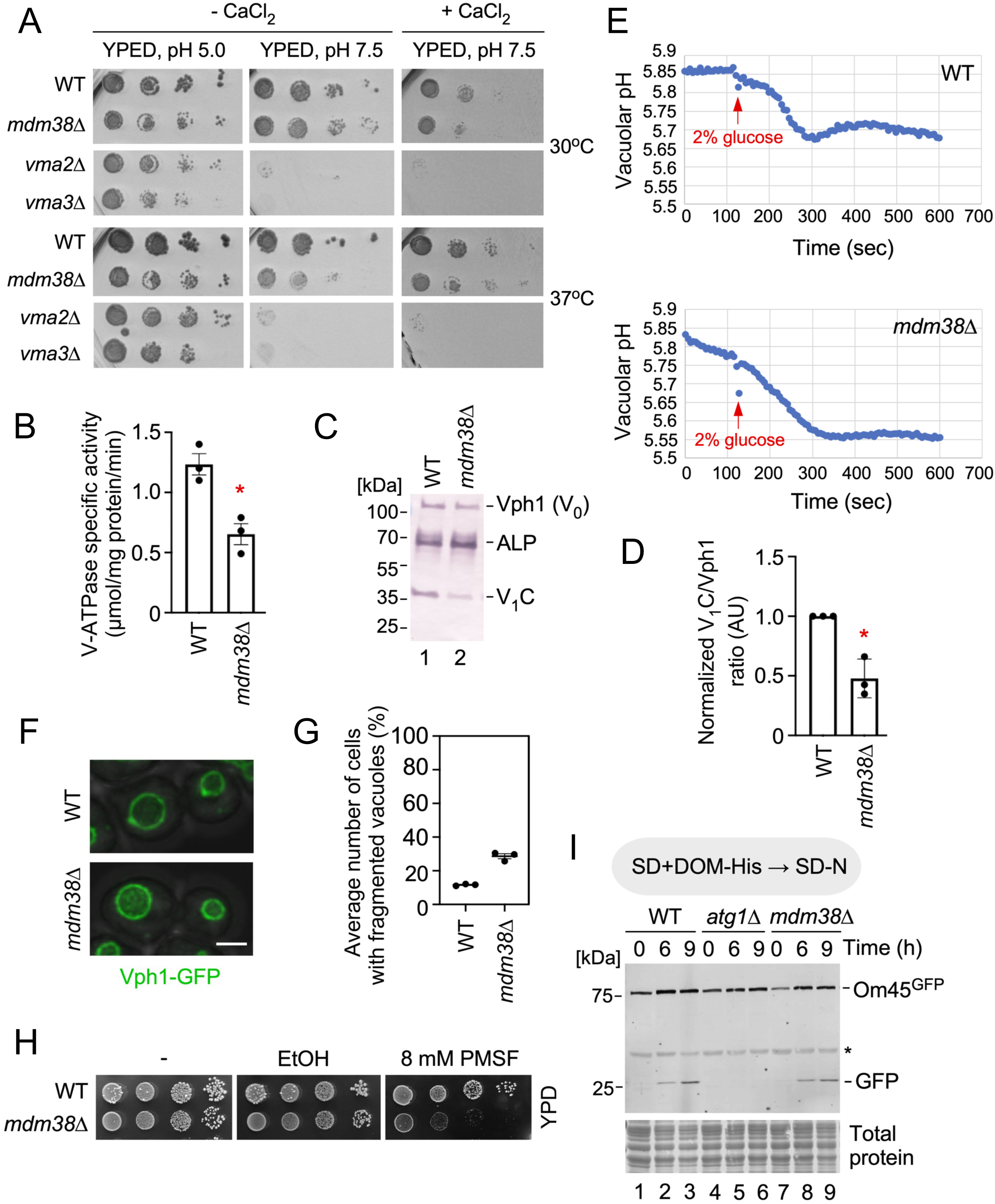
Assessment of vacuolar functions in cells deleted for Mdm38. (A) Growth assays at 30°C and 37°C of indicated yeast strains on yeast extract-peptone-dextrose (YEPD) medium buffered to pH 5 or pH 7.5 ± 60 mM calcium chloride (CaCl2). (B) V-ATPase specific activity in the vacuoles isolated from wild-type or *mdm38*Δ mutant cells. Mean ± SD. for three biological replicates. Asterisk indicates a statistically significant difference by *t*-test (*p<0.05). (C) Representative immunoblot showing steady-state levels of the indicated V-ATPase subunits and the vacuolar membrane protein alkaline phosphatase (ALP, loading control) in vacuoles isolated from wild-type or *mdm38*Δ cells. (D) Quantification of the V-ATPase subunits V1C to Vph1 ratios across three biological replicates. Bars show mean ± SD; n=3 biological replicates. Asterisks indicate a statistically significant difference by *t*-test (*p<0.05). (E) Representative traces of vacuolar pH-sensitive fluorescent dye 2’,7’-Bis-(2-carboxyetyl)-5- (and-6)-carboxyfluorescein acetoxymethyl ester (BCEF) staining of wild-type and *mdm38*Δ cells. Cells were labeled with BCEF, washed, and deprived of glucose for less than 30 min. Fluorescent ratios were monitored starting at time 0. Glucose was added to a final concentration of 2% at 120 sec, and monitoring continued for indicated periods of time. The plots represent a typical response of 3 biological replicates for each strain. (F) Images of wild-type and *mdm38*Δ cells expressing the vacuolar marker Vph1-GFP. Images are max intensity projections of a Z-stack. Scale bar, 2 μm. (G) Quantification of the average percentage of cells containing fragmented vacuoles in the indicated strains. Vacuole fragmentation was defined as any mother cell containing greater than seven Vph1-GFP vacuole lobes. Each dot represents the average of an individual imaging replicate. Bars show mean ± SEM. At least 427 cells were counted per strain. (H) Growth of indicated yeast strains on YPD medium ± ethanol (EtOH) or ethanol-dissolved 8 mM phenylmethylsulfonyl fluoride (PMSF). (I) Immunoblot of the mitochondrial outer membrane protein Om45-GFP, a mitophagy reporter, and free GFP released as a result of Om45-GFP processing in the vacuoles of indicated strains cultured in synthetic medium during 9-hr nitrogen starvation course. The asterisk denotes a non- specific band. Total protein staining was used as a control for protein loading.

While the levels of the membrane-bound Vo sector subunit Vph1 subunit were similar between strains, the V1 sector subunit C was reduced in *mdm38*Δ, suggesting diminished V-ATPase activity may result from impaired assembly. Measurements of vacuolar pH in intact cells using the BCECF probe [47] indicated that, despite a slightly slower response to glucose, the *mdm38*Δ mutant achieved a vacuolar pH comparable to wild-type cells (Fig. 6E).

The *mdm38*Δ strain also displayed a minor (∼28%) increase in vacuolar fragmentation (Fig. 6F, 6G), which was negligible, as suggested by the stable vacuolar pH. The growth of *mdm38*Δ mutant cells was sensitive to high doses of phenylmethylsulfonyl fluoride (PMSF) (Fig. 6H), which inhibits vacuolar protease activity and results in the accumulation of autophagic bodies within the vacuole [48], indicating potentially compromised protease activity within the less acidic vacuole. However, measurements of mitophagy flux revealed no significant disruption in the Mdm38-deficient strain (Fig. 6I).

In summary, while the loss of Mdm38 leads to a modest reduction in V-ATPase activity, and vacuolar morphological changes, these phenotypes likely represent secondary responses to broader cellular perturbations rather than primary effects directly impacting mitochondrial homeostasis.

### Mitochondrial proteases cooperatively link ion and protein homeostasis in the IMM

To explore whether other mitochondrial mutants share similar phenotypic traits with the *mdm38*Δ strain, we conducted a screen using a mini-collection of mutants deleted for non-essential genes encoding for mitochondrial proteome constituents. Each strain was tested for tolerance to cobalt chloride and Conc. A, compounds known to perturb mitochondrial function and protein quality control, relative to WT cells, phenotypes observed in *mdm38*Δ cells. This screen identified three mutant strains, deleted for *PCP1*, *PIM1*, or *YME1,* with growth patterns similar to those of the *mdm38*Δ strain (Fig. 7A). Interestingly, whereas the *yme1*Δ strain exhibited marked resistance to Conc. A, its tolerance to cobalt chloride was less pronounced compared to the *mdm38***Δ** strain, suggesting divergence in the tolerance mechanisms for these two compounds.

**Figure 7.**
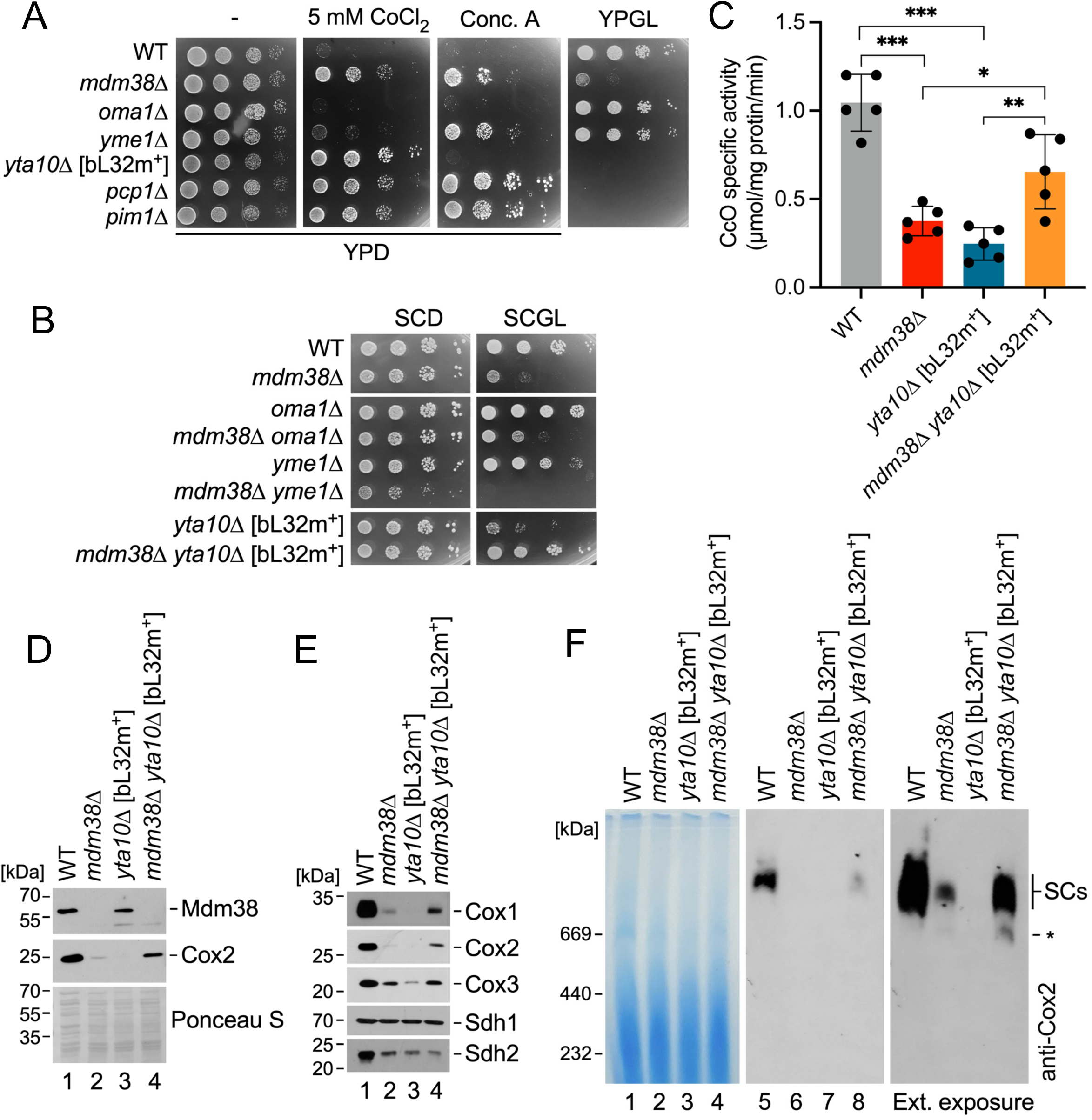
Mitochondrial proteases contribute to *mdm38*Δ-associated phenotypes, which can be mitigated by inactivation of the m-AAA protease. (A) Growth assays of indicated yeast strains on YPD ± 5mM CoCl2 or 100 nM concanamycin A and on YPGL media. Note that the *yta10*Δ strain lacking the m-AAA protease subunit harbors an engineered version of the bL32m mitoribosomal protein to ensure its respiratory competence. (B) Growth assays of indicated yeast strains on SCD and SCGL media. (C) Specific enzymatic activities of complex IV (CcO) in mitochondrial lysates from indicated strains. Bars show mean ± SD; n=5 biological replicates. Asterisks indicate a statistically significant difference by one-way ANOVA test (*p<0.05, **p<0.01, ***p<0.001). (D) Immunoblot showing steady-state levels of Mdm38 and a core CcO subunit Cox2 in the mitochondria from indicated strains. Ponceau S staining was used as a control for protein loading. (E) Western blot analysis showing steady-state levels of indicated proteins in mitochondria from the cells described above. (F) BN-PAGE analysis of respiratory chain supercomplexes (SCs) visualized with antibodies against complex IV core subunit Cox2 in mitochondrial lysates from indicated strains. Coomassie G250-stained membrane shows equal amounts (100 µg) of total mitochondria that were analyzed.

Notably, the genes identified in our screen encode multifaceted mitochondrial quality control factors that – among other roles – have been implicated in the regulation of mitochondrial proteostasis and IMM integrity [21,22]. Therefore, we further explored genetic interactions between Mdm38 and mitochondrial quality control proteases, focusing on IMM-localized enzymes due to the location of Mdm38 within this membrane. Deleting the IMM proteases Yme1, Pcp1, or Atp23 in the *mdm38*Δ strain had no appreciable effect on its respiratory growth, whereas deletion of the Oma1 protease provided only a marginal improvement in respiratory competence (Fig. 7B and data not shown), indicating a limited role of Oma1 in this context.

To test the effect of the m-AAA protease inactivation, we used a modified *yta10*Δ [bL32m^+^] strain in which the mitoribosomal subunit bL32m is engineered to be processed by the mitochondrial processing peptidase instead of the m-AAA protease, allowing mitoribosome assembly and translation in the absence of the Yta10 subunit [23,24]. Remarkably, we found that in contrast to the *mdm38*Δ and *yta10*Δ [bL32m^+^] single mutants, the *mdm38*Δ *yta10*Δ [bL32m^+^] cells were able to propagate on respiratory medium upon prolonged incubation (Fig. 7B). Due to poor metabolic labeling in mitochondria isolated from these older cultures (16h), direct assessment of mitochondrial translation products was challenging (data not shown). However, paralleling the above observation, mitochondria isolated from the 48h-old *mdm38*Δ *yta10*Δ [bL32m^+^] cells showed significantly increased CcO activity and stabilization of CcO subunits and respiratory supercomplexes (Fig. 7C-F), indicating that m-AAA protease plays a major role in IMM protein turnover in Mdm38-deficient mitochondria. Interestingly, in contrast to the CcO subunits, the steady-state levels of SDH subunit Sdh2 remained decreased in the *mdm38*Δ *yta10*Δ [bL32m^+^] strain, suggesting the effect may be specific to IMM-anchored and/or mitochondria-borne proteins. Similarly, Yta10 deletion had no appreciable effect on the *mdm38*Δ mutant’s increased tolerance to cobalt chloride (Fig. S7), indicating that cobalt resistance might operate independently of m-AAA protease function.

Collectively, these findings support a model (Fig. 8) whereby the loss of Mdm38 triggers proteostatic stress in the IMM, activating mitochondrial protein quality control factors such as the m-AAA protease. However, when the m-AAA activity is unrestrained, it disrupts respiratory chain complex assembly and stability, as well as the function of key ion transporters, leading to a range of pleiotropic phenotypes. This model suggests that Mdm38 functions to maintain the delicate balance of IMM proteostasis and ion homeostasis, and its loss unleashes compensatory stress responses that exacerbate mitochondrial dysfunction.

**Figure 8.**
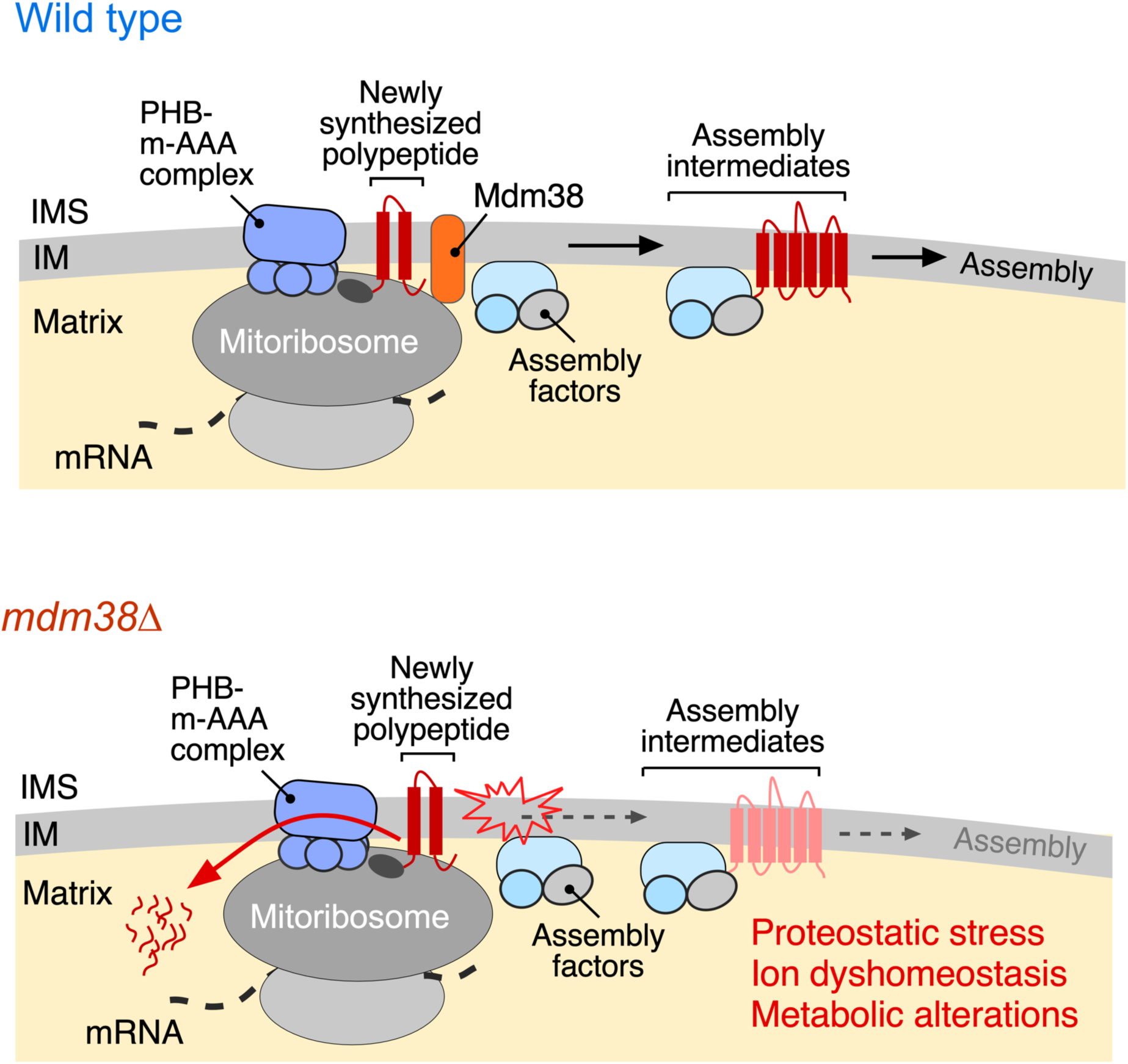
Model depicting the involvement of Mdm38 in the IMM protein and ion homeostasis. The model posits that Mdm38 acts as a molecular steward for newly synthesized polypeptides as they come off the mitoribosome and are integrated into the IMM. Loss of Mdm38 impairs this process causing proteostatic stress and activating IMM quality control proteases such as prohibitin:m-AAA triaging complex. See Discussion for additional details.

## DISCUSSION

The present study unravels a previously unrecognized mechanism that links proteostatic processes to subcellular ion homeostasis and provides new insights into the role of Mdm38/LETM1, a conserved IMM-anchored protein implicated in a variety of seemingly divergent functions such as regulation of mitochondrial translation and ion homeostasis [11,14,19,20]. By combining quantitative proteomics, metabolomics, biochemical experiments, and genetics analyses, we identified primary molecular consequences associated with Mdm38 loss, offering a deeper understanding of the phenotypic changes observed in Mdm38-deficient cells.

We demonstrate that Mdm38 is required to maintain IMM proteostasis, which is crucial for the stability and function of respiratory complexes, as well as for IMM permeability to ions such as iron and metabolites. These findings align with previous reports showing that overexpression of certain carrier proteins can partially rescue the respiratory defect in *mdm38*Δ mutants [38]. While earlier research proposed that Mdm38 acts as a K^+^/H^+^ exchanger based on its influence on IMM ion permeability and mitochondrial swelling, with the ionophore nigericin mitigating these effects [17,19,20], to the best of our knowledge, the direct ion exchange activity of Mdm38 has not been demonstrated. Our current data provide an alternative explanation that does not necessarily contradict previous observations (Fig. 8). Rather than attributing the observed phenotypes to a specific loss of K^+^/H^+^ exchanging activity, we propose that these defects may stem from the deregulation of multiple transporters. In line with this notion is our observation that Mdm38-deleted cells are resistant to high doses of nigericin, which triggers IMM protein quality control mechanisms in wild-type mitochondria.

Furthermore, our data suggest that deregulation of transporters in Mdm38-deficient cells may be selective, as mitochondrial import capacity and thus IMM permeability appear largely unaffected. This observation, coupled with our metabolomics analyses, offer a potential explanation for the previously reported inability of the LETM1-depleted mammalian cells to utilize pyruvate [13]. Indeed, we observed a significant decrease in the mitochondrial pyruvate carrier subunits among other proteins. Even though in our hands the human LETM1 protein showed only marginal ability to rescue the Mdm38-deficient phenotype – likely due to evolutionary divergence and/or the specific genetic background of the *mdm38*Δ strain– our findings, in conjunction with previous studies [49,50], support the notion that the underlying pathways are generally conserved across species.

Mdm38 function is related to mitochondrial translation. Our results are consistent with the earlier findings indicating that, although Mdm38 associates with mitoribosomes, it is not essential for their assembly or IMM tethering [30,31]. However, we show that mitochondrial translation per se is partially impaired in *mdm38*Δ cells. The primary defect appears to lie in the stability of newly synthesized polypeptides, which are destabilized due to unregulated activity of the m-AAA protease and potentially other proteases. This scenario is reminiscent of what has been described in mutants lacking the CcO assembly factor Coa2, wherein newly synthesized Cox1 is rapidly degraded by the stress-activated Oma1 protease [28,51].

Although structural information on Mdm38 remains limited, a previous study identified a 14-3-3-like receptor domain in Mdm38, suggesting its involvement in protein-protein associations with newly synthesized polytopic proteins, such as Cox1 and complex III Cyt.*b* subunit [14,30]. It is tempting to speculate that Mdm38 functions as a molecular steward for nascent polypeptides as they emerge from the mitoribosome and are integrated into the IMM. Conversely, Mdm38 loss of function disrupts this process, leading to proteostatic stress and activation of IMM quality control proteases. Indeed, m-AAA inactivation allows for a slow, but adequate assembly of the OXPHOS complexes in the *mdm38*Δ cells. Our findings are consistent with a recent report that describes a prohibitin:m-AAA protease molecular triage system located in the vicinity of the mitoribosomal tunnel exit, where it determines the fate of newly synthesized mitochondrial OXPHOS subunits [10]. Notably, that report identified Mdm38 alongside Oma1 and Yme1 proteases, in high-throughput proximity labeling and chemical crosslinking experiments targeting the proteins associated with the ribosomal tunnel exit and the prohibitins.

Further supporting our model, prior studies have shown that newly synthesized polypeptides co-purify with Mdm38 and the mitoribosomes [11,14,30], although it remains to be determined whether or not Mdm38’s association with nascent polypeptides is direct. Furthermore, this model is supported by our finding that Mdm38-deleted cells exhibit signatures of a known proteostatic response, resembling the mitochondrial precursor overaccumulation stress response [34] and are sensitive to expression of mito-DHFR constructs that challenge IMM proteostasis.

Additionally, the original report by Rep and Grivell identified Mdm38’s partner protein Mba1 as a multicopy genetic suppressor in strains devoid of the m-AAA subunit Yta10 [52], further supporting the idea that Mdm38 and its interactors play a crucial role in IMM proteostasis.

Our study also unravels a previously unrecognized link between IMM proteostatic mechanisms and subcellular iron metabolism. We show that one of the primary consequences of Mdm38 loss is disrupted iron homeostasis, which impinges on mitochondria-to-vacuole communication. Such disruption likely contributes to some of the secondary phenotypes previously reported for the *mdm38*Δ mutant. Importantly, altered iron homeostasis is a shared trait among several mutants lacking mitochondrial quality control factors such as the i-AAA protease Yme1, mitochondrial rhomboid protease Pcp1, and LON protease Pim1. This discovery provides an important step toward a deeper understanding of how these quality control factors contribute to maintaining the functional integrity of both mitochondria and vacuoles, and may also shed light onto the mechanisms underpinning the etiology of genetic and aging-associated disorders associated with mitochondrial and lysosomal dysfunction. In this context, our results align with recent findings by Hughes et al. [53], which linked age-related decline in vacuolar function and perturbed iron homeostasis to progressive decline in mitochondrial health. Our data suggest further reciprocity between mitochondrial and vacuolar homeostatic mechanisms and indicate that they are crucial for cellular physiology. The emerging connection between mitochondrial function and vacuolar iron regulation opens new avenues for understanding how these organelles collaborate in maintaining cellular health.

Given the relevance of these findings to disease, it will be important to validate our findings in the context of relevant disease models, including those of Wolf-Hirschhorn syndrome [49,54,55] and neurodegenerative disorders such as Parkinson’s and Alzheimer’s diseases, where mitochondrial and lysosomal dysfunction are key pathological features [56,57].

## MATERIALS AND METHODS

### Yeast strains, plasmids, and culturing conditions

Yeast strains used in the present work are listed in Supplementary Table S1. Strains made for this study were generated by *in vivo* homologous recombination using PCR-amplified pAG60 plasmid-derived deletion cassettes targeting the gene of interest and containing either *KanMX4* antibiotic resistance or various prototrophic markers. Mitochondrial matrix targeted DsRed and TagBFP were expressed from pRS305-mito-DsRed [58] and pRS305-ADH1::mito-TagBFP (this study) integrated at the *LEU2* locus. Vph1, Vps39, and Mdm12 were tagged at the endogenous loci using plasmids pKT128 pFA6a–link–yEGFP–SpHIS5 [59] and pFA6-yoHalo::HIS [60]. The complete list of plasmids used in this study is provided in the Supplementary Table S2. All constructs were sequence verified using whole-plasmid nanopore DNA sequencing.

Depending on the experiment, cells were cultured in yeast extract-peptone (YP) or synthetic complete (SC) medium supplemented with relevant amino acids [61] and containing 2% glucose, galactose, or 2% glycerol/D-lactic acid mix as a sole carbon source. Unless indicated otherwise, cultures of interest were grown overnight, serially diluted, and tested for growth as before [25,62].

### Mitochondrial isolation and assays

Intact mitochondria-enriched fractions were isolated from yeast cells using established protocols [63]. When necessary, mitochondrial fractions were further purified using discontinuous Histodenz gradients as described before [64]. Total mitochondrial concentrations were determined using the Coomassie Plus assay kit (Thermo Scientific). Mitochondria were snap-frozen in liquid nitrogen and stored at -80 °C.

The specific activities of mitochondrial enzymes cytochrome *c* oxidase, aconitase, and succinate dehydrogenase were measured according to established protocols [65,66].

### In organello labeling of mitochondrial translation products

Metabolic labeling of mitochondrial translation products in isolated mitochondria with ^35^S-methionine in isolated yeast mitochondria was carried out according to published protocols [67].

### Phospholipid analysis

Overnight cultures of indicated strains grown in YPD at 30°C were diluted to an OD600 = 0.2 in 2 mL of YPD supplemented with 1 μCi/mL ^14^C-Acetate and grown shaking at 240 rpm for ∼24 hr in a 30°C water bath. Yeast were collected at 1690 x g for 5 min, washed with 2 mL distilled water, and re-centrifuged at 1690 x g for 5 min. Following resuspension of each yeast pellet with 0.3 mL MTE buffer (0.65M mannitol, 20mM Tris, pH 8.0, and 1 mM EDTA) containing 1mM PMSF, 10 μM leupeptin, and 2 μM pepstatin A, the resulting slurries were transferred to1.5 mL microcentrifuge tubes containing ∼0.1 mL glass beads, and each tube sealed with parafilm. Yeast were mechanically obliterated by vortexing on high for ∼30 min at 4 °C. The resulting extract, separated from the glass beads and unbroken yeast following a 2 min 250 × g centrifugation at 4°C, was further centrifuged for 5 min at 13,000 × g at 4°C to yield crude mitochondria. Following liquid scintillation, equal amounts of labeled crude mitochondria were transferred to 5 mL borosilicate tubes containing1.5 mL 2:1 chloroform:methanol and vortexed for 30 min on medium-high on a laboratory bench. To stimulate phase separation, 0.3 mL 0.9% (w/v) NaCl was added to each sample prior to a 1 min vortex on high. Following a 1000 rpm spin in a clinical centrifuge for 5 min at room temperature, the upper aqueous phase was aspirated into the radioactive waste, and the remaining organic phase washed with 0.25 mL 1:1 Methanol:H2O, vortexed on high for 30 seconds at room temperature, and centrifuged as before. The lower organic phase was transferred to a fresh tube and dried down under a stream of nitrogen. The dried lipid extracts were resuspended in 13 μL of chloroform and loaded on SILGUR-25 (Machery-Nagel) TLC plates that had been washed with chloroform, pretreated with 1.8% (w/v) boric acid in 100% ethanol, and activated at 95°C for at least 30 min. Phospholipids were resolved using chloroform:ethanol:H2O:triethylamine (30:35:7:35), after which the plates were air-dried in a fume hood, wrapped in saran wrap and developed using a K-screen and FX-Imager (Bio-Rad Laboratories).

### Protein import into isolated mitochondria

Precursor proteins of interest were synthesized *in vitro* using the pGEM4Z-SP6p-Su9-DHFR plasmid [68], rabbit reticulocyte lysate (Promega) and Express^35^S protein labeling mix (Perkin Elmer) and imported into isolated mitochondria according to established protocols [69]. Briefly, mitochondria isolated from cells of interest were incubated with radiolabeled protein precursors in the presence of energizing mix (2 mM NADH, 2mM ATP and an ATP-regenerating system consisting of 2.5 mM malate, 2.5 mM succinate, 1 mM creatine phosphate, and 0.1 mg/mL creatine kinase) to obtain a highly energized state of mitochondria and power the import reaction. Remaining unimported precursor protein was removed by addition of 100 µg/mL proteinase K and incubation for 30 min on ice; the reaction was stopped by addition of 2 mM PMSF.

#### Mitophagy flux assessment

Cells were pre-grown for one day in 2 ml of SD+DOM-His medium [70] and inoculated into 50 mL of SD+DOM-His at OD600 = 0.005 for approximately 16 h until they reached OD600 = 0.5-1. Then, 25 ODs of cells were washed twice with sterile deionized water and resuspended in 25 mL of SD-N medium at OD600 = 1 for 0, 6, 9 h time-course. Sampling of SD-N cultures and western blot analyses were performed as previously described [70].

#### Vacuolar isolation and assays

Vacuolar vesicles were isolated by flotation on Ficoll gradients as described [71]. ATPase activity was assessed in a coupled enzyme assay; activity sensitive to 200 nM Concanamycin A is designated V-ATPase activity. In order to assess the levels of V-ATPase subunits in vacuolar vesicles, 5 µg of vesicle protein was solubilized and separated by SDS-PAGE, then transferred to nitrocellulose. The blot was probed with monoclonal antibodies 10D7 against Vph1 (Vo subunit a) and 7A2 against the V1C subunit [47], and monoclonal antibody 1D3A10 against alkaline phosphatase (Invitrogen). To quantitate relative subunit levels, blots from three separate vacuole preparations were scanned, and peak intensities were determined using the plot profile feature in FIJI. The ratio of V1C to Vph1 signal was calculated for both wild-type and *mdm38*Δ, and then the mutant ratio was normalized to the wild-type. Vacuolar acidification was determined using BCECF as described [47]. Briefly, cells were grown to log-phase in fully supplemented minimal medium, then labeled with BCECF-AM. After washing to remove free dye, cells were maintained on ice in the absence of glucose for 15-30 min. Fluorescence was then continuously measured at excitation wavelengths 450 and 490 nM and emission wavelength 535 nm at 30°C. Fluorescence ratios at 450 and 490 nm were calculated for each time point and converted to pH using a calibration curve determined for each cell culture [47].

#### Confocal fluorescence microscopy

For imaging, strains were grown to mid-log phase (OD600 of 0.7-1.0) in synthetic complete + 2% wt/vol dextrose media with 2x adenine (SCD) at pH 6.4. Cells were concentrated by centrifugation and imaged on a 4% wt/vol agarose pad on a depression slide.

Imaging was performed using a Leica SP8 confocal microscope equipped with HyD detectors and a 63x 1.4NA oil immersion objective. All images were deconvolved using Huygens Essential (Scientific Volume Imaging) prior to analysis.

HaloTag labeling was performed as previously described [72]. Briefly, JFX650 HaloTag ligand [73] was added from a 1 mM stock in DMSO to a log-phase yeast culture in SCD pH 6.4 media to a final concentration of 1 μM, and labeling was performed for 30 min at 24°C with shaking. Excess dye was removed by filter washing. Washed cells were resuspended in SCD pH 6.4 and imaged.

All image analysis was performed using Fiji [74]. To quantify vacuole morphology, individual Z-stacks were max intensity projected and manually scored for the total number of vacuole lobes per cell. Vacuole fragmentation was defined as any mother cell containing greater than seven Vph1-positive vacuole lobes [75]. At least 427 cells were counted per background over three imaging replicates.

To quantify the number and intensity of Mdm12 ERMES foci, Z-stacks were average projected, and a background fluorescence value, obtained from regions of the image that did not contain cells, was subtracted. The Mdm12 channel was isolated, the fluorescence signal was smoothed using a Gaussian blur filter, and individual foci were segmented using the “Analyze Particles” tool in Fiji. The segmented ROIs were then used to quantify the signal of individual Mdm12 foci from the original average projections. To be included in the quantification, foci needed to be larger than 4 pixels and above the background fluorescence value. At least 132 cells and 685 Mdm12 foci were analyzed over three imaging replicates for each genetic background.

#### Omics and data analysis

Quantitative multidimensional protein identification (MudPIT) analysis was carried out essentially as described [76]. Fifty million WT or *mdm38*Δ mid-log cells or 100 µg of mitochondria-enriched fractions from the respective strains were used.

Gas chromatography-mass spectrometry (GC-MS) metabolomics analysis was carried out at the University of Utah Center for Iron and Heme Disorders Metabolomics Core. Fifty million cells per sample were mixed with boiling 75% ethanol solution containing the internal standard d4-succinic acid (Sigma, 293075). Boiling samples were vortexed and incubated at 90 °C for 5 min. Samples were then incubated at -20 °C for 1 hr. After incubation, the samples were centrifuged at 5,000 x *g* for 10 minutes at 4 °C. The supernatant was then transferred from each sample tube into a labeled, fresh 13x100mm glass culture tube. A second standard was then added d27-myristic acid (CDN Isotopes, D-1711). Pooled quality control samples were made by removing a fraction of the collected supernatant from each sample, and process blanks were made using only extraction solvent and no cells. The samples were then dried *en vacuo*. GC-MS analysis was performed with an Agilent 7200 GC-QTOF fit with an Agilent 7693A automatic liquid sampler. Dried samples were suspended in 40 µL of a 40 mg/mL O-methoxylamine hydrochloride (MOX, MP Bio, 155405) in dry pyridine (EMD Millipore, PX2012-7) and incubated for one hour at 37 °C in a sand bath. Twenty-five µL of this solution was added to auto-sampler vials. 60 µL of N-methyl-N-trimethylsilyltrifluoracetamide (MSTFA with 1%TMCS, Thermo, TS48913) was added automatically via the auto sampler and incubated for 30 minutes at 37 °C. Following incubation, samples were vortexed, and 1 µL of the prepared sample was injected into the gas chromatograph inlet in the split mode with the inlet temperature held at 250 °C. A 10:1 split ratio was used for analysis for the majority of metabolites. Any metabolites that saturated the instrument at the 10:1 split were analyzed at a 50:1 split ratio. The gas chromatograph had an initial temperature of 60 °C for one minute, followed by a 10 °C/min ramp to 325 °C and a hold time of 10 minutes. A 30-meter Agilent Zorbax DB-5MS with 10 m Duraguard capillary column was employed for chromatographic separation. Helium was used as the carrier gas at a rate of 1 mL/min. Below is a description of the two-step derivatization process used to convert non-volatile metabolites to a volatile form amenable to GC-MS. Data was collected using MassHunter software (Agilent). Metabolites were identified, and their peak area was recorded using MassHunter Quant. Metabolite identity was established using a combination of an in-house metabolite library developed using pure purchased standards, the NIST library, and the Fiehn library. Data were analyzed using in-house software to prepare for analysis by the MetaboAnalystR software tool [77].

Inductively coupled plasma mass spectrometry (ICP-MS) analyses of transition metals content in whole cells or gradient-purified mitochondria were carried out using an established protocol [78].

#### Electrophoretic techniques and immunoblotting

Blue-native (BN)-PAGE analysis of mitochondrial protein complexes solubilized with 1% digitonin was performed as described [62] using 5-13% gradient polyacrylamide gels. Individual proteins were separated by denaturing SDS-PAGE using 10% polyacrylamide gels. Separated proteins or protein complexes were transferred to nitrocellulose or PVDF membranes, respectively, and incubated with primary antibodies of interest and goat anti-mouse and anti-rabbit HRP-coupled secondary antibodies (sc-2005 and sc- 2030, Santa Cruz Biotechnology). Protein bands were then visualized by incubation of membranes with chemiluminescence reagents (Thermo Scientific) and visualized on X-ray films (Thomas Scientific). The primary antibodies used in this study are listed in the Supplementary Table S3.

#### Statistical methods

At least three biological replicates per sample were obtained for each represented experiment. Statistical analyses were carried out using Microsoft Excel or GraphPad Prism 10.2 software. Unless specified otherwise, data are shown as means ± standard deviation (S.D.). Statistical significance was determined by the one-way ANOVA with post-hoc Tukey’s comparison test and considered significant at *p*≤0.05.

## SUPPLEMENTAL INFORMATION

Supplemental information can be found online.

## ACKNOWLEDGEMENTS

We thank Rabab Mahdi and Drs. Ganapathi Kandsamy, and Jonathan Dietz for early investigations related to this study. We thank Drs. Dennis Winge and Diane Ward (University of Utah); Alexander Tzagoloff (Columbia University); Nikolaus Pfanner (University of Freiburg); Johannes Herrmann (University of Kaiserslautern); Peter Rehling (University of Gottingen); Dejana Mokranjac (Ludwig-Maximilians University); Valeria Culotta (Johns Hopkins University); Luke Lavis (HHMI/Janelia Research Campus); Charles Boone (University of Toronto); and Roland Lill (University of Marburg) for reagents. We are grateful to Dr. Javier Seravalli and the University of Nebraska Redox Biology Center Biophysics Core for help with ICP-MS analyses and the members of the University of Utah Center for Iron and Heme Disorders Metabolomics Core2 for their help with metabolomics analyses. We thank Drs. Adam Hughes (University of Utah), Martin Ott (University of Gothenburg), Charles Boone (University of Toronto), and members of the Khalimonchuk lab for fruitful discussions and critical reading of the manuscript.

## AUTHOR CONTRIBUTIONS

Conceptualization, I.B., P.M.K., A.B., and O.K.; Methodology, I.B., G.M.S., S.F.A., E.J.B., E.M.G., A.M., J.A.W., J.C.C., M.A.R., T.Y.N., M.T., S.M.C., and A.B.; Formal analysis, I.B., G.M.S., E.M.G., A.M., J.A.W., J.C.C., T.Y.N., T.S., L.L.L., S.M.C., P.M.K., A.B., and O.K.; Investigation, I.B., G.M.S., E.M.G., J.A.W., J.C.C., M.A.R., T.S., L.L.L., S.M.C., P.M.K., A.B., and O.K.; Writing -Original draft, I.B., J.C.C., T.Y.N., L.L.L., S.M.C., P.M.K., and O.K.; Visualization, I.B., J.C.C., T.Y.N., L.L.L., S.M.C., P.M.K., A.B., and O.K.; All authors reviewed and edited the manuscript. Supervision, T.Y.N., T.S., L.L.L., P.M.K., and O.K.; Project administration, L.L.L., P.M.K., A.B., and O.K.; Funding acquisition, S.M.C., L.L.L., P.M.K., and O.K.

## Supplemental Information

**Table S1.**
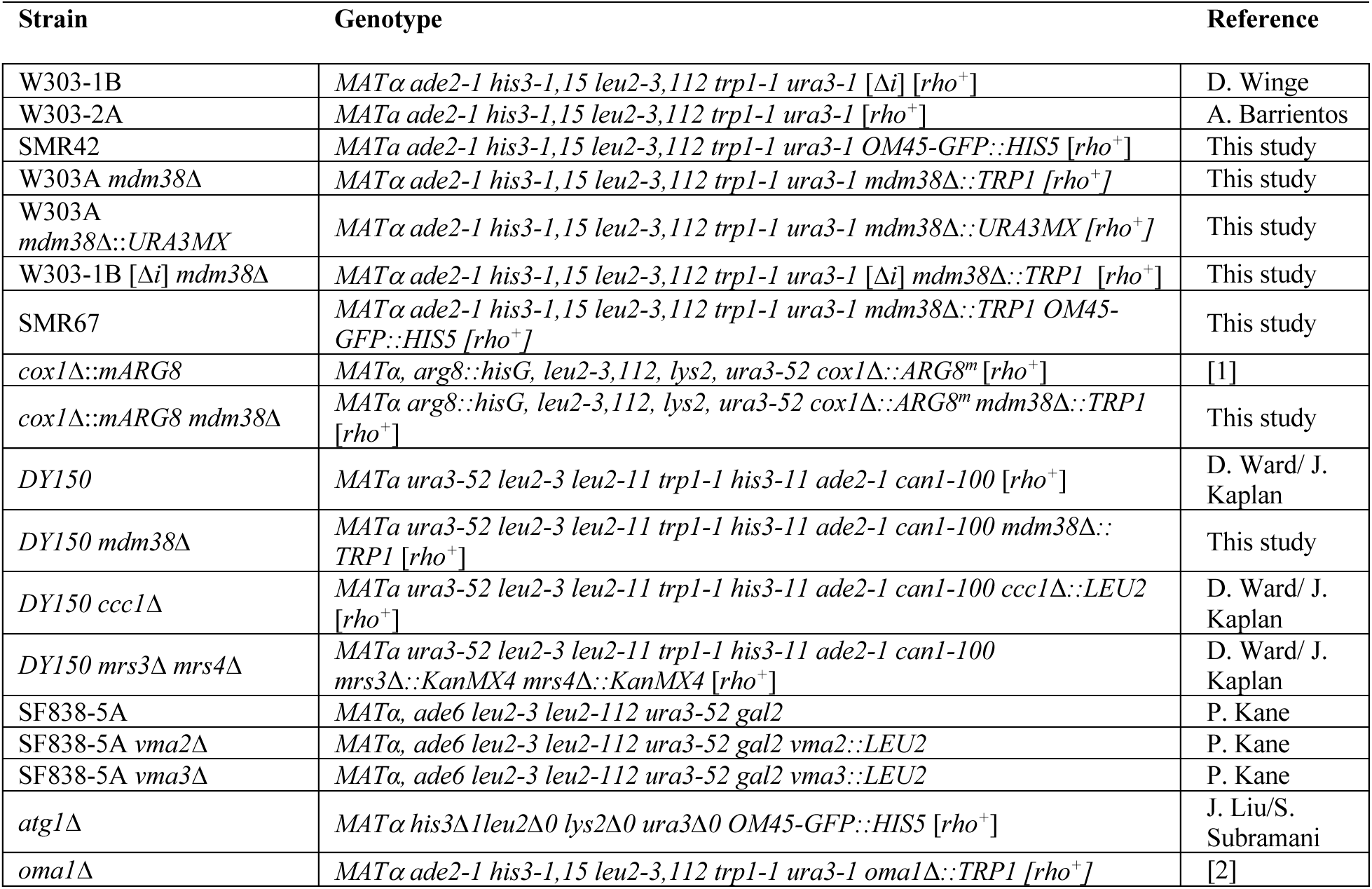
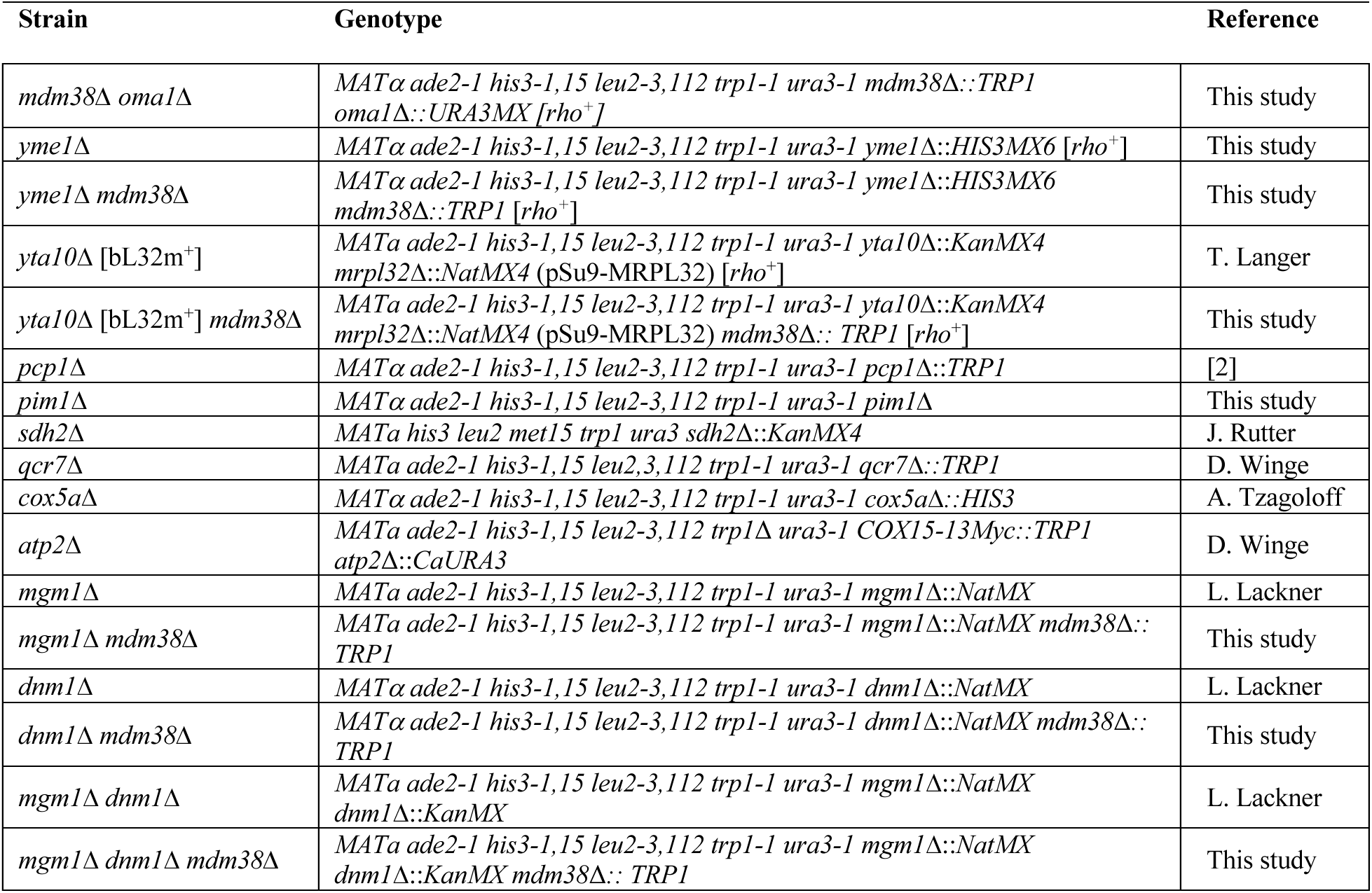
Yeast strains used in this study.

**Table S2.** Plasmids used in this study.

| Plasmid | Genotype | Reference |
| --- | --- | --- |
| Oma1-13Myc | pRS425-Oma1-Myc-13×Myc | [2]4 |
| pFA6a-Trp1 | Used for gene deletions | [3] |
| pFA6a-His3MX6 | Used for gene deletions | [3] |
| pAG60-Ura3MX | Used for gene deletions | [4] |
| *MDM38* | pRS416-Mdm38 | [5] |
| *LETM1* | PR426-ADH1<sub>pr</sub>-LETM1-3HA-CYC1<sub>tem</sub> | [6] |
| *IMS-DHFR* | pYES2- Cyb<sub>2</sub>(1-107)-DHFR | [7] |
| *mx-DHFR* | pYES2- Cyb<sub>2</sub>(1-107, Δ19)-DHFR | [7] |
| *Mrs3-FLAG* | pCM185-Mrs3-FLAG | [8] |
| FET3pr-LacZ |  | D. Ward |
| pKT128 pFA6a-link–yEGFP–SpHIS5 | Used for Vph1 and Mdm12 endogenous tagging | [9] |
| pFA6-yoHalo::HIS3 | Used for Vps39 endogenous tagging | [10] |
| *Mito-BFP* | pRS305-ADH1::mito-TagBFP | This study |
| *Mito-Red* | pRS305-mito-DsRed | [11] |
| *Oxa1-Ura3* | pRS313-Oxa1-Ura3 | [12] |
| *pGAL-MRS3-MRS4* |  | C. Boone |

**Table S3.** Primary antibodies used in this study.

| Antibody | Dilution | Secondary | Type/Source |
| --- | --- | --- | --- |
| c-Myc | 1:1000 | Mouse | Monoclonal, Roche (11667149001) |
| Porin | 1:5000 | Mouse | Monoclonal, Invitrogen (11667149001) |
| Mdm38 | 1:2000 | Rabbit | Serum (gift to T. Barrientos from J. Hermann's lab) |
| Mdm38 | 1:5000 | Rabbit | A gift from N. Pfanner |
| LETM1 | 1:5000 | Rabbit | Polyclonal, Proteintech (16024-1-AP) |
| Aco1 | 1:5000 | Rabbit | A gift from R. Lill |
| Aac2 | 1:10000 | Rabbit | A gift from C. Koehler |
| Kar2 | 1:10000 | Rabbit | Polyclonal, Santa Cruz Biotechnology (Y-115; sc-33630) |
| Cox1 | 1:1000 | Mouse | Monoclonal, Abcam (ab110270) |
| Cox2 | 1:2000 | Mouse | Monoclonal, Abcam (ab110271) |
| Cox3 | 1:2000 | Mouse | Monoclonal, Abcam (ab110259) |
| Sdh1 | 1:2000 | Rabbit | A gift from D. Winge |
| Sdh2 | 1:2000 | Rabbit | A gift from D. Winge |
| Rip1 | 1:2000 | Mouse | A gift from B. Trumpower |
| Sod2 | 1:500 | Rabbit | Serum, a gift from V. Culotta |
| GFP | 1:2000 | Mouse | Roche Diagnostics (11814460001) |
| DHFR | 1:5000 | Mouse | R&D Systems (MAB7934) |
| Mrs3 | 1:500 | Rabbit | Serum, a gift from D. Ward |
| Lipoic acid | 1:5000 | Rabbit | Polyclonal, Abcam (ab58724) |

## Supplemental Figure Legends

**Figure S1.**
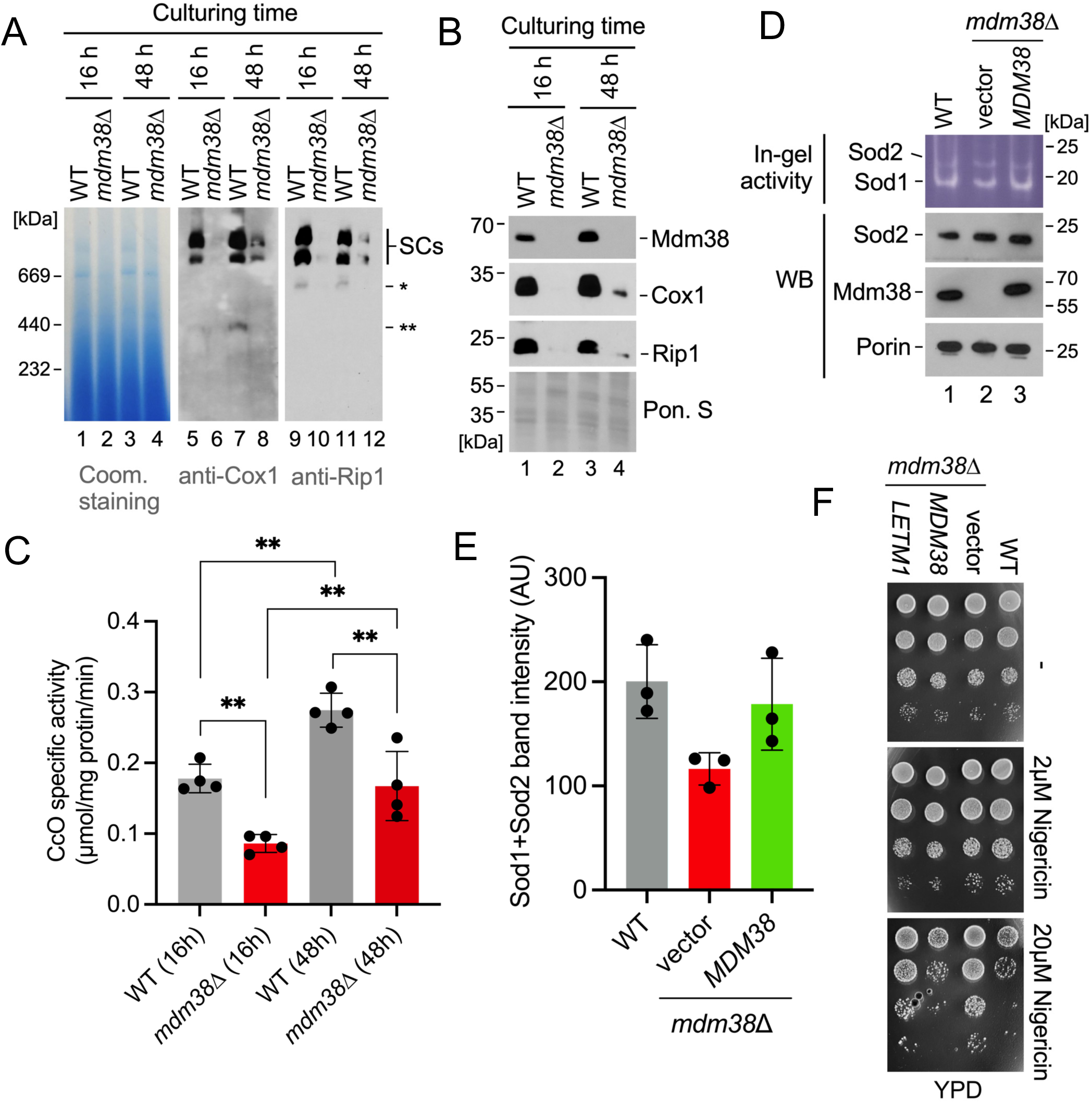
Related to Figure 1 Unique growth and respiratory phenotypes of Mdm38- deleted cells. (A) BN-PAGE analysis of respiratory chain complex III-complex IV supercomplexes (SCs) visualized with antibodies against complex IV core subunit Cox1 in mitochondrial lysates from wild-type and *mdm38*Δ cells cultured in the non-repressive galactose-containing medium for 16 or 48 h. (B) Steady-state levels of indicated proteins in mitochondrial lysates from panel A. (C) Specific enzymatic activities of complex IV (CcO) in mitochondrial lysates from cells described above. Bars show mean ± SD; n=5 biological replicates. Asterisks indicate a statistically significant difference by *t*-test (*p<0.05, **p<0.01, ***p<0.001). (D) Top panel: Representative in-gel analysis of superoxide dismutase activity following native gel electrophoresis of the mitochondria from indicated cells. Bottom panel: Immunoblot analysis of steady-state Sod2, Mdm38, and porin levels in the mitochondria from respective cells. (E) Quantification of Sod1 and Sod2 in-gel staining signals. Bars are mean ± SD; n=3 biological replicates. (F) Growth of indicated yeast cells on YPD medium ± 2 or 20 µM nigericin.

**Figure S2.**
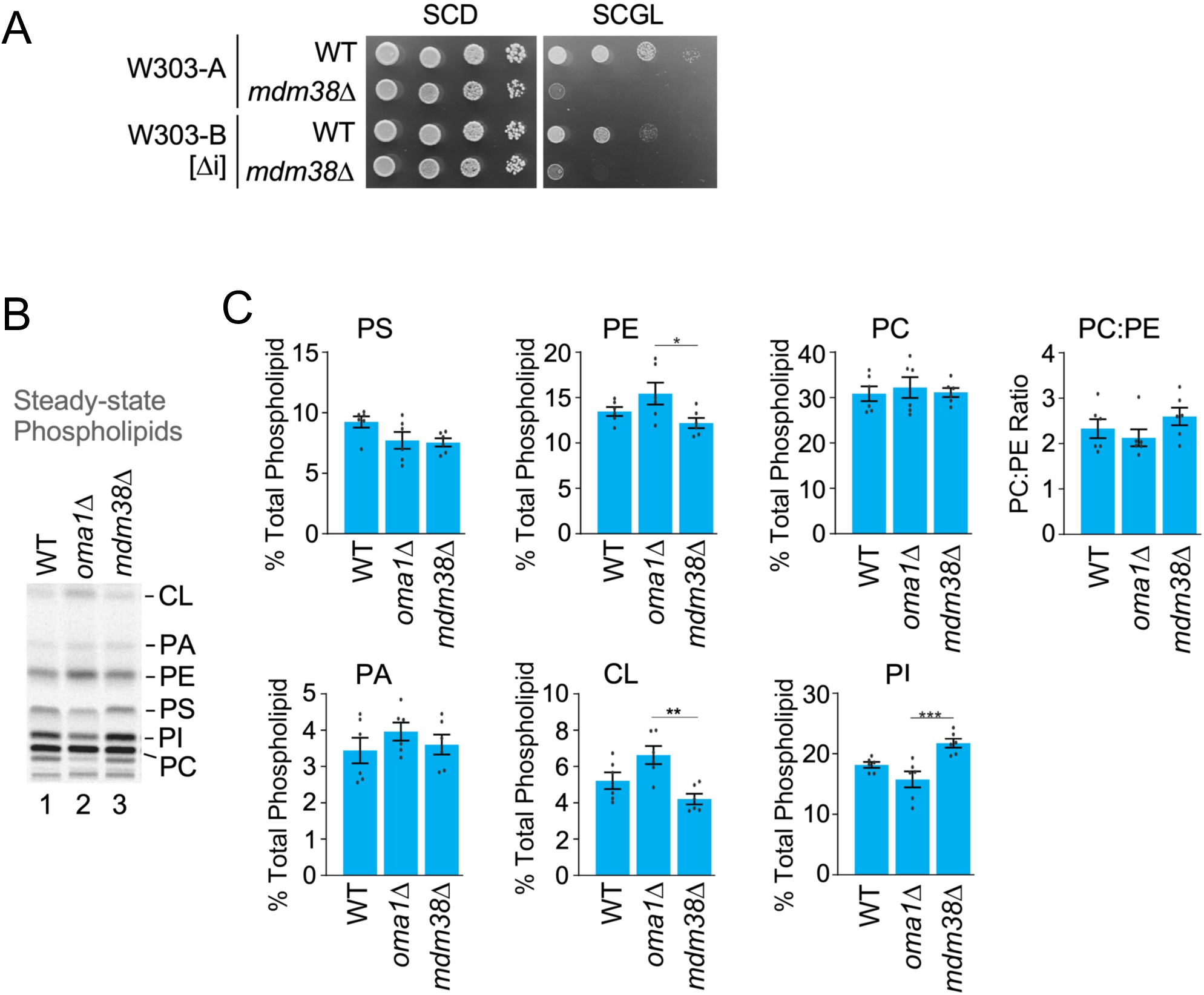
Related to Figure 2. Mitochondrial DNA introns or altered phospholipid composition do not contribute to phenotypes upon Mdm38 deletion. (A) Growth of indicated yeast strains on SCD or SCGL medium. *[Δi],* strains devoid of introns in their mitochondrial DNA. (B) Analysis of phospholipid content in mitochondria from indicated cells. Representative image of thin layer chromatography (TLC)-separated, ^14^C-Acetate-labeled phospholipids extracted from mitochondria and visualized with autoradiography. PC, phosphatidylcholine; PI, phosphatidylinositol; PS, phosphatidylserine; PE, phosphatidylethanolamine; PA, phosphatidic acid; CL, cardiolipin. (C) Quantification of indicated phospholipid species from the TLC analysis above. Bars are mean ± SEM; n=3 biological replicates. Asterisks indicate a statistically significant difference by *t*-test (*p<0.05, **p<0.01, ***p<0.001).

**Figure S3.**
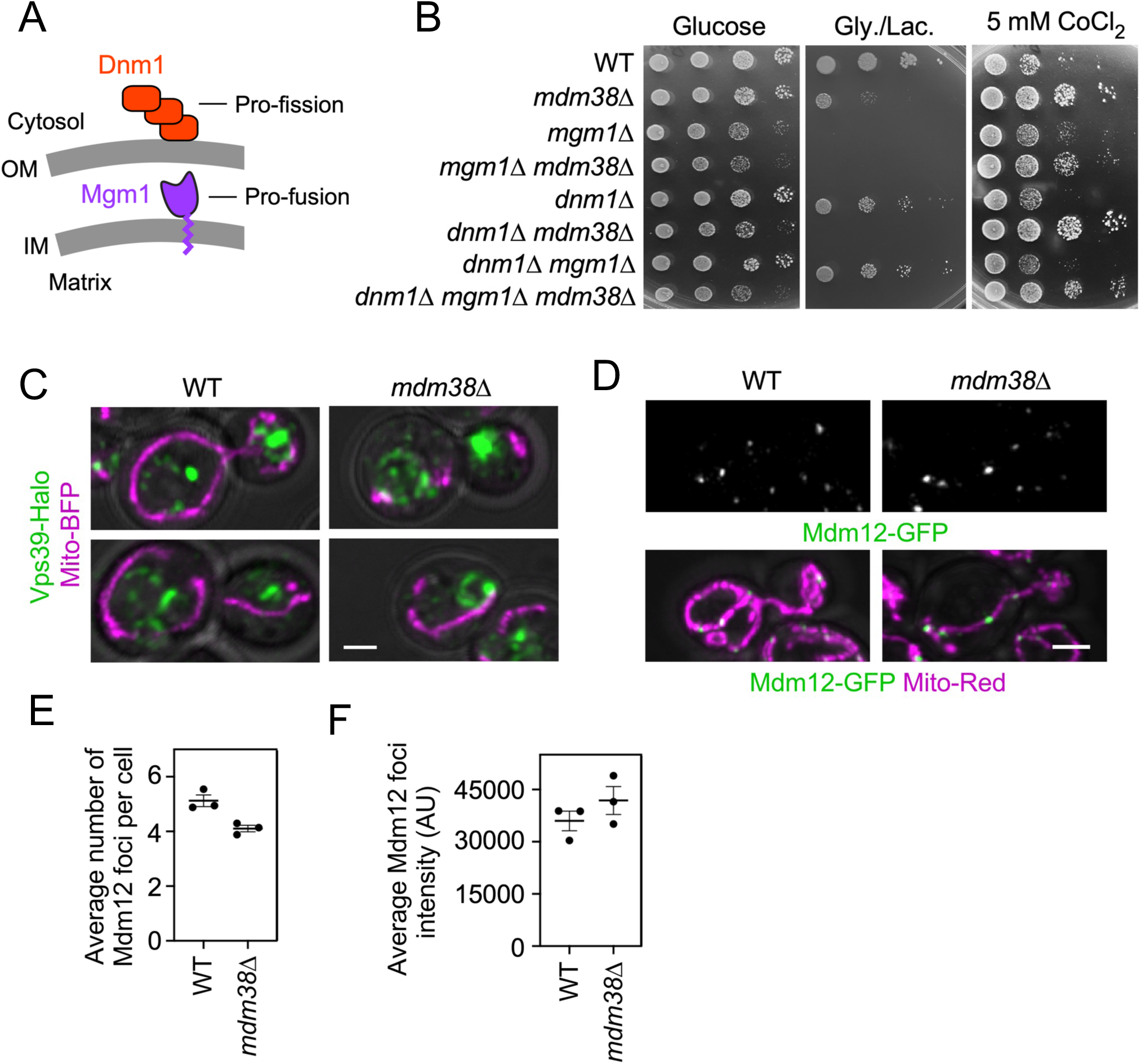
Related to Figure 2. Impaired mitochondrial network morphology does not drive respiratory deficient phenotype in the *mdm38*Δ mutant. (A) Schematic depiction of the key factors regulating mitochondrial network behavior in yeast. Dnm1 is pro-fission factor, and Mgm1 is a pro-fusion protein. (B) Growth of indicated yeast strains on YPD ± 5 mM cobalt chloride (CoCl2) or YPGL medium. (C) Representative images of a mitochondrial marker, MitoBFP, with the vCLAMP component Vps39 under endogenous expression levels. In both wild-type and *mdm38*Δ cells, the proximity between Vps39-positive structures and mitochondria can be seen, which likely represents vCLAMPs. Images are maximal intensity projections of two central slices of a Z-stack. Scale bar, 2 μm. (D) Representative images of wild-type and *mdm38*Δ cells expressing Mdm12-GFP, an ER-mitochondria encounter structure (ERMES) component, and the mitochondria marker MitoRed. Scale bar, 2 μm. (E) Quantification of the number of ERMES foci in the indicated genetic backgrounds. Each dot represents the average of an individual imaging replicate, and the error bars represent the SEM between the three replicates. At least 132 cells were counted per background. (F) Quantification of the average ERMES foci intensity in the indicated genetic backgrounds. Each dot represents the average of an individual imaging replicate. Bars are mean ± SEM; n=3 biological replicates. At least 685 foci were counted per background.

**Figure S4.**
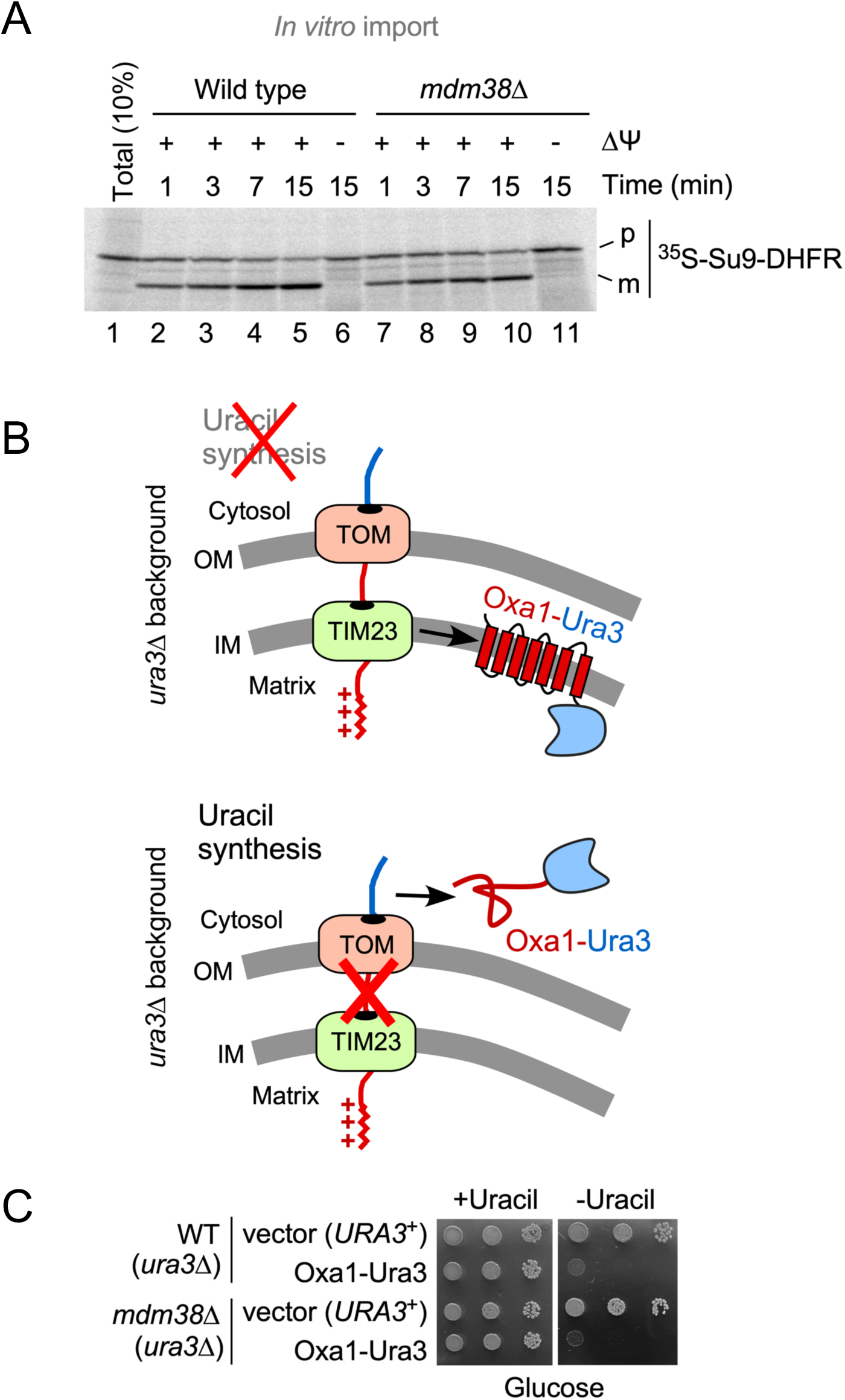
Related to Figure 2. Mitochondrial protein import is unaffected by the Mdm38 deletion. (A) Autoradiogram analysis of the radiolabeled Su9-DHFR model substrate imported *in vitro* into energized (+ΔΨ) or depolarized (-ΔΨ) mitochondria isolated from wild-type and *mdm38*Δ cells for indicated periods. p, non-imported precursor Su9-DHFR; m, mature imported form of Su9-DHFR. (B) Schematic depiction of the Oxa1-Ura3 genetic reporter used to monitor mitochondrial protein import *in vivo*. (C) Growth assay of indicated cells on SCD medium ± uracil.

**Figure S5.**
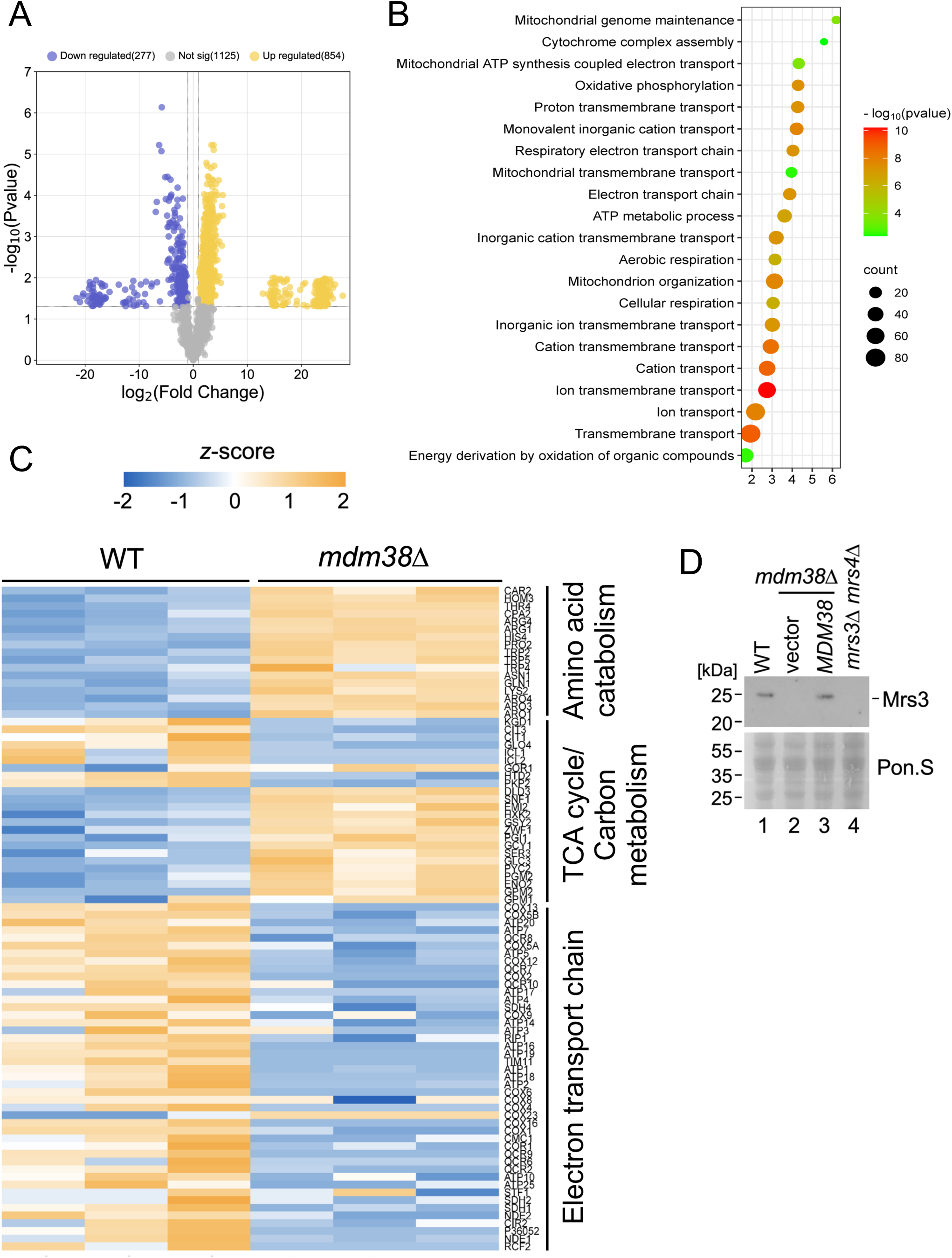
Related to Figure 3. Proteomic changes in Mdm38 deleted cells. (A) Volcano plot showing proteomic changes in wild-type and *mdm38*Δ cells. The blue circles depict downregulated proteins in the *mdm38*Δ mutant with FDR< 0.05. (B) Protein set enrichment analysis comparing wild-type and *mdm38*Δ cells. (C) Enlarged version of the heat map presented in Figure 3C to highlight changes in individual proteins in identified mitochondrial processes clustered groups. The heat map shows the abundance of proteins from proteomic analysis of mitochondria from log-phase wild-type and *mdm38*Δ cells. Blue, downregulated; yellow, upregulated. (D) Western blot analysis of steady-state levels of Mrs3 in indicated strains. Ponceau S staining was used as a control for protein loading.

**Figure S6.**
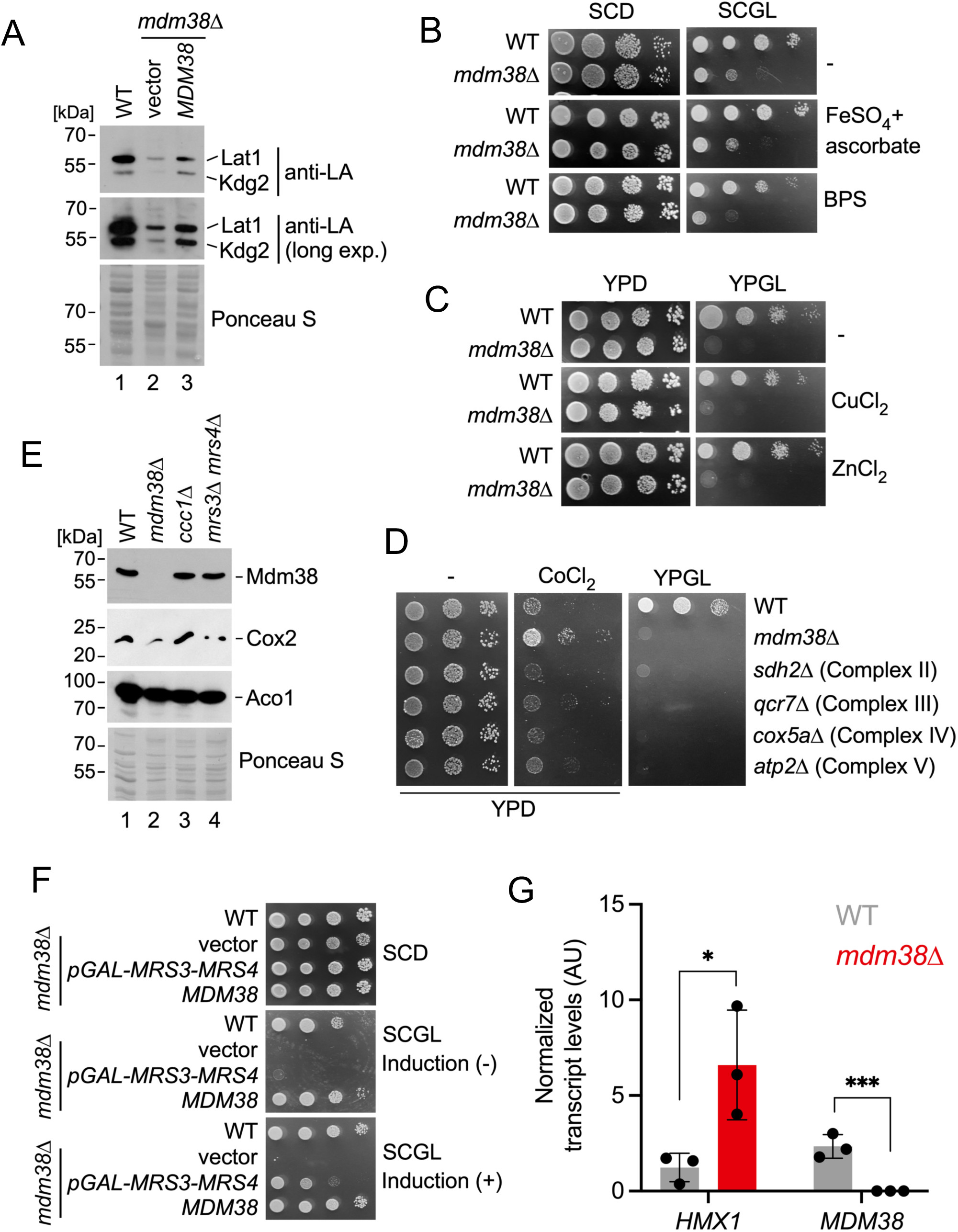
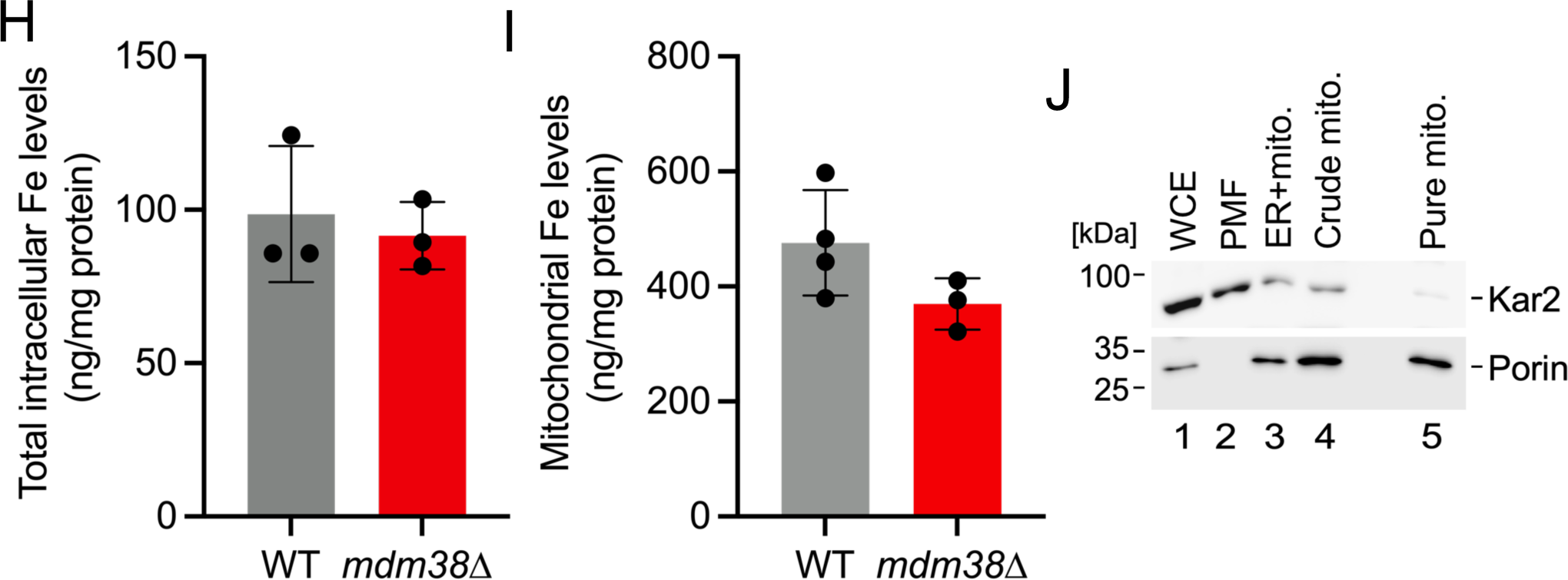
Related to Figure 5. Assessment of iron uptake in the Mdm38-deleted cells. (A) Western blot analysis of lypoilated mitochondrial proteins, Lat1 and Kdg2, visualized by antibodies against lipoic acid (LA) moiety. Ponceau S staining was used as a control for protein loading. (B) Growth assay of indicated cells on SCD and SCGL medium ± bathophenanthrolinedisulfonate (BPS) or ferrous ammonium sulfate (Fe). (C) Growth assay of indicated strains on YPD or YPGL medium ± indicated divalent metal salts. (D) Growth of indicated yeast strains on YPD medium ± 5 mM cobalt chloride (CoCl2) or on respiratory YPGL medium. (E) Western blot analysis of steady-state levels of Mdm38, Cox2, and aconitase in indicated strains. The proteins were visualized with respective antibodies. Ponceau S staining was used as a control for protein loading. (F) Induced concomitant expression of *MRS3* and *MRS4* partially suppresses growth defects in the *mdm38*Δ mutant. Growth assay of indicated cells on SCD and SCGL medium. (G) Levels of the indicated transcripts in the WT and *mdm38*Δ cells. Bars are mean ± SD; n=3 biological replicates. (H) Total intracellular iron levels in wild-type and *mdm38*Δ cells analyzed by ICP-MS. Bars are mean ± SD; n=3 biological replicates. (I) Mitochondrial iron levels tend towards decline but do not reach statistical significance in the organelles isolated from the *mdm38*Δ mutant. Purified mitochondrial fractions from wild-type and *mdm38*Δ cells were analyzed for iron content by ICP-MS. Bars are mean ± SD; n=3-4 biological replicates. (J) Representative immunoblot showing purity of histodenz gradient-purified mitochondrial fractions analyzed in panel I. WCE, whole-cell extract; PMF, post-mitochondrial fraction. Porin is a mitochondrial marker, Kar2 = ER. Note that the residual Kar2 signal in the mitochondrial fraction is due to its presence in the mitochondria-associated membranes (MAMs).

**Figure S7.**
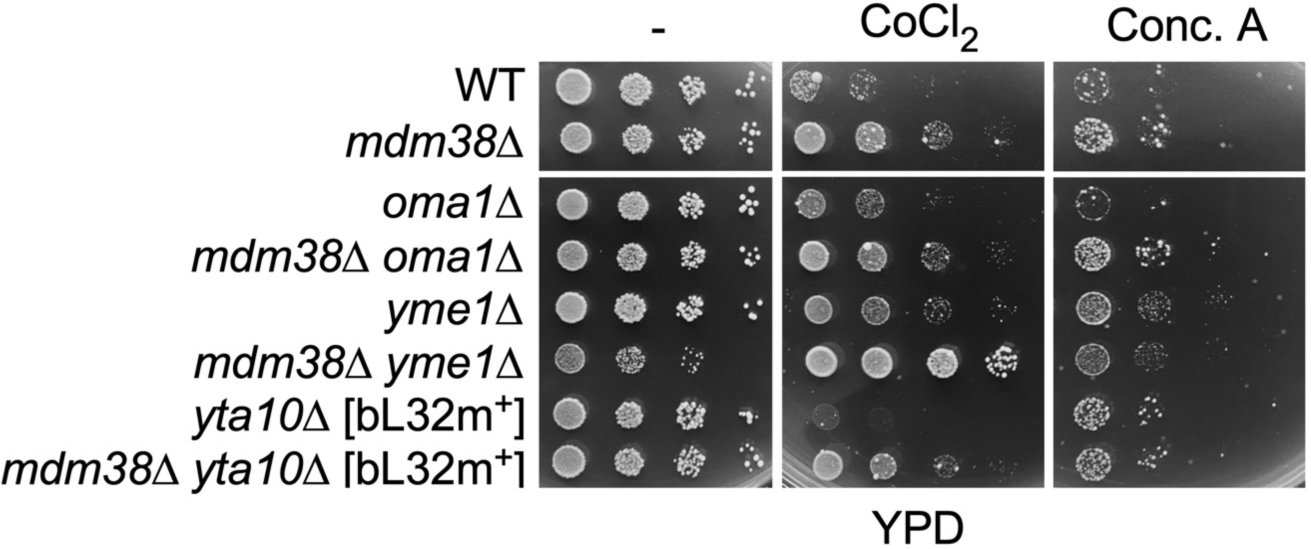
Related to Figure 7. Synthetic genetic analysis of *MDM38* and genes encoding key IMM proteases. Growth assay of indicated strains on YPD medium ± 5 mM cobalt chloride (CoCl2) or 100 nM concanamycin A.

